# ARAP2 regulates responses to interferon-gamma by restricting SOCS1

**DOI:** 10.1101/2025.04.28.650977

**Authors:** Narelle Keating, Karen Doggett, Grace M Bidgood, Lizeth G Meza Guzman, Laura F Dagley, Kunlun Li, Anna Gabrielyan, Carolina Alvarado, Bailey E Williams, Benjamin J Broomfield, Brigette C Duckworth, Colin Hockings, Jumana Yousef, Evelyn Leong, Rhiannon Morris, Andrew Kueh, Alexandra L Garnham, Jean-Laurent Casanova, Stephanie Boisson-Dupuis, Jeffrey J Babon, Edmond M Linossi, Michelle D Tate, Joanna R Groom, Sandra E Nicholson

## Abstract

Interferon-gamma (IFNγ) is critical for immunity against intra-macrophagic pathogens, signaling through the JAK-STAT pathway to induce a tyrosine-phosphorylation cascade that ensures a potent immune response. Excessive JAK-STAT signaling can drive hyperinflammation and autoimmunity, and thus signaling is tightly and selectively regulated by the IFNγ-inducible protein, Suppressor of Cytokine Signaling 1 (SOCS1). SOCS1 inhibits signaling by directly blocking JAK kinase activity. Here we identified a SOCS1-interacting partner, ARAP2 that fine-tunes SOCS1 function. We report that tyrosine 415 in ARAP2 binds the SOCS1-Src Homology 2 (SH2) domain and limits the ability of SOCS1 to inhibit IFNγ signaling. Our findings show that ARAP2 promotes the IFNγ response through a phosphorylation dependent interaction with the negative regulator SOCS1.

## INTRODUCTION

The cytokine interferon-gamma (IFNγ; or type II IFN) is indispensable for host defense and orchestrating immunity. Originally identified as a macrophagic-activating factor (MAF), loss of IFNγ function due to the presence of neutralizing autoantibodies or to genetic deficiencies that affect the production of or response to IFNγ, has severe consequences in human disease^1^. Individuals with rare monogenic inborn errors in IFNγ immunity are susceptible to infectious pathogens, in particular intra-macrophagic mycobacterial or fungal infections, associated with diseases such as Mendelian Susceptibility to Mycobacterial Disease (MSMD)^2^ and Tuberculosis (TB)^3^. The critical role of IFNγ in immunity is supported by extensive genetic studies in mice, with deletion of either *IFNγ* or the IFNγ receptor subunits (*IFNGR1* or IFNGR1), resulting in increased susceptibility to tuberculous^4^ and non-tuberculous^5^ mycobacteria, Gram-negative^6^ and some Gram-positive^7^ bacteria, several protozoa^8^ and fungi^9^, and some viruses, such as SARS-CoV-2^10^ and Influenza A^11^.

Although there are no known type II interferonopathies, there are inherited conditions that result in enhanced IFNγ, such as genetic errors in *SOCS1* (Suppressor Of Cytokine Signaling 1), the critical intracellular negative regulator of IFN signaling^12, 13, 14, 15^. In mice, genetic deletion of *Socs1* is lethal, due to excessive IFNγ signaling and hyperinflammation^16^. Neonatal death can be rescued by genetic deletion of *Ifng*^16^, or its downstream signal transducer and activator of transcription (*Stat)1*^17^. In humans, inborn errors of *SOCS1* have been reported in patients presenting with an array of inflammatory phenotypes, including immune dysregulation with multi-system autoimmunity, autoimmune diseases such as SLE and chronic autoimmune cytopenia, and malignancy^12, 18, 19, 20, 21, 22, 23^. Notably, cells derived from individuals with loss-of-function SOCS1 variants consistently displayed elevated IFNγ signaling^12^, underscoring the importance of precisely regulating the IFNγ response.

IFNγ initiates signaling by binding to its cognate receptors, IFNGR1 and IFNGR2 at the cell surface^24, 25^, leading to transactivation of the Janus kinases (JAK), JAK1 and JAK2^26^. The JAKs phosphorylate tyrosine residues in the cytoplasmic domain of IFNGR1^24, 26^, enabling recruitment and phosphorylation of STAT1 protein dimers^27, 28, 29, 30^. Phosphorylation of STAT1 results in a change in dimer conformation, facilitating translocation of active STAT1 homodimers to the nucleus to bind IFNγ-activated site (GAS) elements^31, 32, 33^ and reprogram the ‘interferome’ by regulating transcription of interferon stimulated genes (ISG)s^34^. SOCS1 is a critical ISG induced by IFNγ^16^ (and by type I^35^ and III^36^ IFNs), that inhibits IFNγ signaling in a classic negative feedback loop to prevent hyperactivation and excessive inflammation.

The SOCS family (including CIS and SOCS1-7) recognize their targets through a central Src homology 2 (SH2) domain that selectively binds linear phosphotyrosine (pTyr) motifs within the target proteins^37^. The family share a conserved “SOCS box” motif, which recruits an E3 ubiquitin ligase complex to mediate the proteasomal degradation of SH2-bound proteins^38, 39^. Additionally, SOCS1 and SOCS3 interact with the JAKs via a non-canonical SOCS-SH2 interaction with a GQM motif on JAK1, JAK2 and TYK2, but not JAK3, allowing the SOCS kinase inhibitory region (KIR) to directly and potently inhibit JAK enzymatic activity by blocking access to the JAK substrate binding groove^40, 41, 42^. Notably, mutation of the SOCS1-SH2 domain (*mus musculus*; SOCS1 R105A, *homo sapiens*; SOCS1 R104A) or the SOCS1-KIR (*mus musculus*; SOCS1 F59A, *homo sapiens*; SOCS1 F58A) renders SOCS1 unable to inhibit IFNγ signaling^43^.

Our study aimed to discover proteins that might regulate SOCS1 control of IFNγ signaling. We identified and characterized a novel SOCS1-SH2 binding partner, the GTPase activating protein, ARAP2, and described a role for ARAP2 in promoting IFNγ signaling in cell lines and during *in vivo* influenza infection. Here, we report that ARAP2 restricts the ability of SOCS1 to inhibit JAK1 and thus IFNγ signaling, via a high-affinity interaction between phosphorylated tyrosine 415 in ARAP2 (pY415) and the SOCS1-SH2 domain. We have uncovered a novel role for ARAP2 in influenza control, which suggests that dampening the effects of the pro-inflammatory cytokine IFNγ during severe influenza infection, for instance with IFNγ-neutralizing antibodies, Jakinibs or SOCS1 agonists, could be beneficial in reducing lung injury and morbidity.

## RESULTS

### ARAP2 is a novel, high affinity SOCS binding partner

The identification of SOCS-SH2 domain targets has helped define their *in vivo* function. As part of a campaign to identify new SOCS SH2 domain binders and regulators, we utilized recombinant GST-human (h)CIS^35–173, 202–258^ in complex with hElongin B^1–118^ and hElongin C^17–112^ (GST-CIS/BC) to enrich interacting proteins from cell lysates. Natural killer cells (KYHG-1) were pre-treated with IL-2 to activate JAK-STAT signaling, and the proteasome inhibitor MG132, to reduce degradation of potential SH2-SOCS box targets. Cell lysates were then incubated with GST alone or GST-CIS/BC, prior to analysis of associated proteins by mass spectrometry. In addition to known components of the native CIS E3 ligase complex, such as Cullin 5 and Ring Box protein 2 (RBX2), a number of proteins were significantly enriched by GST-CIS relative to the GST control (**Figure 1A, Supplementary Table 1**). One of the highly enriched proteins identified was ArfGAP with RhoGAP domain, Ankyrin repeat and Pleckstrin-homology domain 2 (ARAP2).

**Figure 1:**
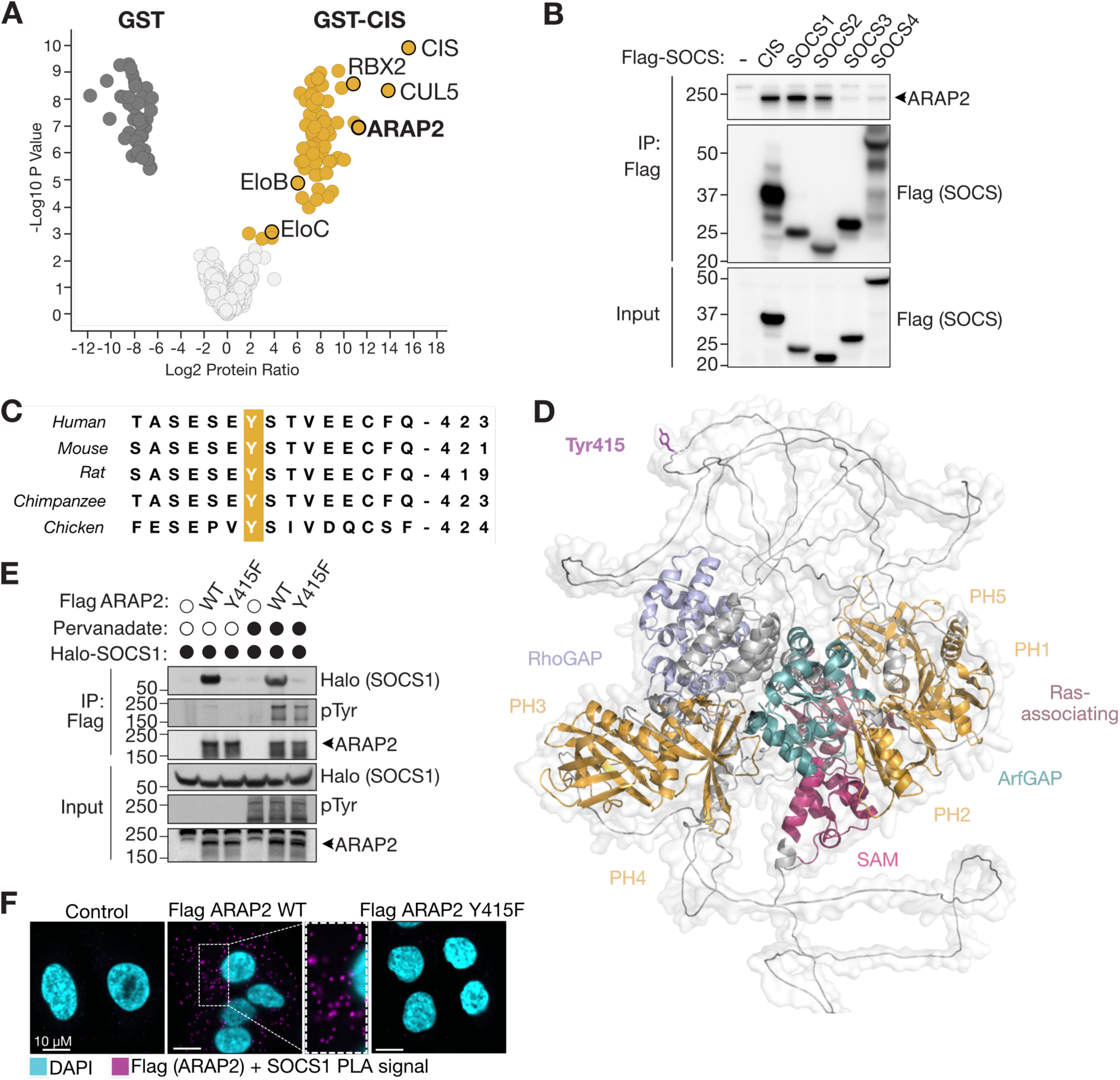
SOCS1 interacts with Tyr415 in ARAP2. **A:** Volcano plot comparing enrichment of GST-CIS bound complexes relative to GST alone. Proteins in yellow indicate at least 2-fold enrichment with GST-CIS compared with GST alone. Proteins in dark grey indicate at least 2-fold enrichment with GST alone compared with GST-CIS. Proteins in light grey were not enriched differently between GST and GST-CIS. **B:** HEK293T cells were transfected with a panel of Flag-tagged SOCS constructs. Flag-SOCS proteins were immunoprecipitated and the presence of endogenous ARAP2 analyzed by SDS-PAGE and immunoblot. **C:** Sequence alignment of ARAP2-Tyr415 and surrounding residues from mammalian species. **D:** AlphaFold2.0 predicted structure of ARAP2 with domains and Tyr415 annotated. **E:** HEK293T cells stably expressing a doxycycline-inducible Halo-SOCS1 construct were transiently transfected with empty vector, Flag-ARAP2 (WT) or Flag-ARAP2 Y415F, and treated with doxycycline overnight to induce SOCS1 expression. Where indicated, cells were treated with pervanadate (PV) for 30 min prior to lysis. Flag-ARAP2 was enriched using anti-Flag antibodies and enrichment of Halo-SOCS1 analyzed by SDS-PAGE and immunoblot. **F:** Proximity ligation assay (PLA). HEK293T cells stably expressing a doxycycline-inducible HaloTag - 3x Flag ARAP2 WT and Y415F construct were treated with doxycycline overnight to induce ARAP2 expression, followed by IFNγ (100 ng/mL) for 6 h to induce SOCS1 expression. Cells were fixed and stained with anti-SOCS1 and anti-Flag antibodies, and DAPI dye (cyan) for confocal analysis using the Zeiss LSM 880 microscope. Pink denotes PLA signal and indicates interaction between SOCS1 and ARAP2. p = phosphorylated, *non-specific band. **A-B**, **E-F:** Representative of three independent experiments. See Supplementary Figure 1 for repeat experiments.

To investigate whether ARAP2 was enriched by other SOCS family members, HEK293T cells were transiently transfected with constructs encoding full-length Flag-tagged CIS, SOCS1, SOCS2, SOCS3 or SOCS4. Flag-SOCS proteins were immunoprecipitated and immunoblotted for ARAP2 enrichment. ARAP2 was present in Flag-CIS, Flag-SOCS1 and Flag-SOCS2 but not Flag-SOCS3 or Flag-SOCS4 immunoprecipitates (**Figure 1B, Supplementary** Figure 1A **& B**). These data suggested that ARAP2 preferentially interacted with the closely-related SOCS family members, CIS, SOCS1 and SOCS2.

We hypothesized that this interaction was mediated by a direct-binding event between pTyr motifs in ARAP2 and the SOCS-SH2 domain. ARAP2 contains 44 Tyr residues, 14 of which were known to be phosphorylated as per the PhosphositePlus database (http://www.phosphositeplus.com). Three of the 14 pTyr sites conformed to a SOCS-SH2 binding consensus sequence (a hydrophobic residue in the +3 position relative to the key pTyr) and were selected for *in vitro* binding studies. Competitive surface plasmon resonance (SPR) was used to determine the relative binding affinities between recombinant SOCS proteins and synthetic peptides derived from ARAP2-pTyr motifs. The highest affinity interaction occurred between SOCS1/BC and the ARAP2-pY415 peptide (IC_50_ = 0.05 µM). CIS/BC also interacted with the pY415 peptide with high affinity (IC_50_= 0.2 µM). In addition, SOCS1 and CIS interacted with pY593, albeit with lower affinity than pY415 **(Table 1, Supplementary** Figure 1J). The affinity between SOCS2 and ARAP2-pY415 peptides (IC_50_= 1.66 µM) was within the observed range for SH2 but was respectively 8- and 30-fold weaker than observed for CIS and SOCS1. The interaction between SOCS3 and all ARAP2 pTyr peptides tested was negligible.

**Table 1:** SPR analysis of SOCS-SH2 binding to ARAP2 phosphopeptides.

| pY | Sequence |  |  |  |  |  |  |  |  |  |  | IC<sub>50</sub> (mM)* |  |  |  |
| --- | --- | --- | --- | --- | --- | --- | --- | --- | --- | --- | --- | --- | --- | --- | --- |
|  | -4 | -3 | -2 | -1 | 0 | +1 | +2 | +3 | +4 | +5 | +6 | CIS | SOCS1 | SOCS2 | SOCS3 |
| **415** | S | E | S | E | pY | S | T | V | E | E | C | 0.2±0.01 | 0.05±0.01 | 1.66±0.11 | 7.52±0.03 |
| **593** | E | K | C | G | pY | L | E | L | R | G | Y | 2.63±0.02 | 3.71±0.02 | 16.35±0.06 | 46.2±0.15 |
| **77** | D | I | P | I | pY | A | N | B | H | K | T | 23.59±0.04 | 35.04±0.04 | 25.88±0.07 | 8.96±0.08 |

\*Mean ± S.D from three independent experiments. pY=phosphorylated tyrosine. SPR curves are shown in Supplementary Figure 1J.

### ARAP2-Y415 is required for interaction with SOCS1

The binding affinity of the SOCS1-SH2 for the ARAP2-pY415 peptide was comparable to peptides derived from the JAK1 activation loop (KD = 0.1-0.6 µM)^41^, indicating that ARAP2-pY415 may represent a physiologically relevant binding site. Tyr415 and the surrounding residues are conserved in mammals, suggesting it is important for ARAP2 structure and/or function **(Figure 1C)**. Furthermore, Tyr415 resides in a region of ARAP2 that is predicted to be unstructured (AlphaFold 2.0^44^; **Figure 1D)**, suggesting it is accessible for SH2-domain binding.

HEK293T cells stably transduced with an inducible Halo-tagged SOCS1 construct (Halo-SOCS1) were transiently transfected to co-express Flag-tagged wild-type (WT) ARAP2 or ARAP2 with Tyr415 mutated to Phe (Flag-ARAP2-WT or Flag-ARAP2-Y415F). Cells were treated with doxycycline to induce Halo-SOCS1 expression and pervanadate (a global tyrosine-phosphatase inhibitor) to maximize tyrosine phosphorylation prior to Flag-ARAP2 enrichment. Halo-SOCS1 interacted with Flag-ARAP2-WT but not Flag-ARAP2-Y415F, in both unstimulated and pervanadate-treated cells **(Figure 1E, Supplementary** Figure 1C **& D)**. pTyr bands corresponding in size to ARAP2 were detected in Flag-ARAP2-Y415F immunoprecipitates with pervanadate treatment, indicating that despite additional pTyr sites being available in ARAP2, SOCS1 selectively interacted with ARAP2-pY415 **(Figure 1E)**. Similarly, ARAP2-Y415 was required for the interaction between ARAP2 and CIS **(Supplementary** Figure 1E-G). Furthermore, a proximity ligation assay was used to confirm a direct interaction between SOCS1 and Flag-ARAP2-WT, but not Flag-ARAP2-Y415F, in the cytoplasm of intact A549 cells **(Figure 1F, Supplementary** Figure 1H **& I)**. Taken together, these data provided evidence for a selective and high-affinity interaction between the SOCS1-SH2 domain and ARAP2-pY415.

### ARAP2-deficient cells have diminished IFNγ signaling

Given that SOCS1 is the key regulator of IFNγ signaling, we investigated whether ARAP2 had a role in IFNγ-induced JAK-STAT signaling. Endogenous ARAP2 was initially depleted using siRNA in a human lung epithelial cell line (A549 cells). *ARAP2*-depletion resulted in reduced STAT1 phosphorylation, compared with cells transfected with scramble siRNA **(Figure 2A, Supplementary** Figure 2A **& B)**, suggesting that ARAP2 was required to promote IFNγ signaling. Once phosphorylated, STAT1 undergoes a conformational change and accumulates in the nucleus to regulate gene transcription^41^. As expected, in control cells transfected with scramble siRNA, STAT1 diffused across the cytoplasm and nucleus under steady-state conditions and rapidly accumulated in the nucleus in response to 30 min IFNγ treatment. Consistent with the reduced STAT1 phosphorylation, *ARAP2* depletion resulted in a two-fold decrease in STAT1 nuclear localization that was evident prior to IFNγ treatment **(Figure 2B, Supplementary** Figure 2C **& D)**.

**Figure 2:**
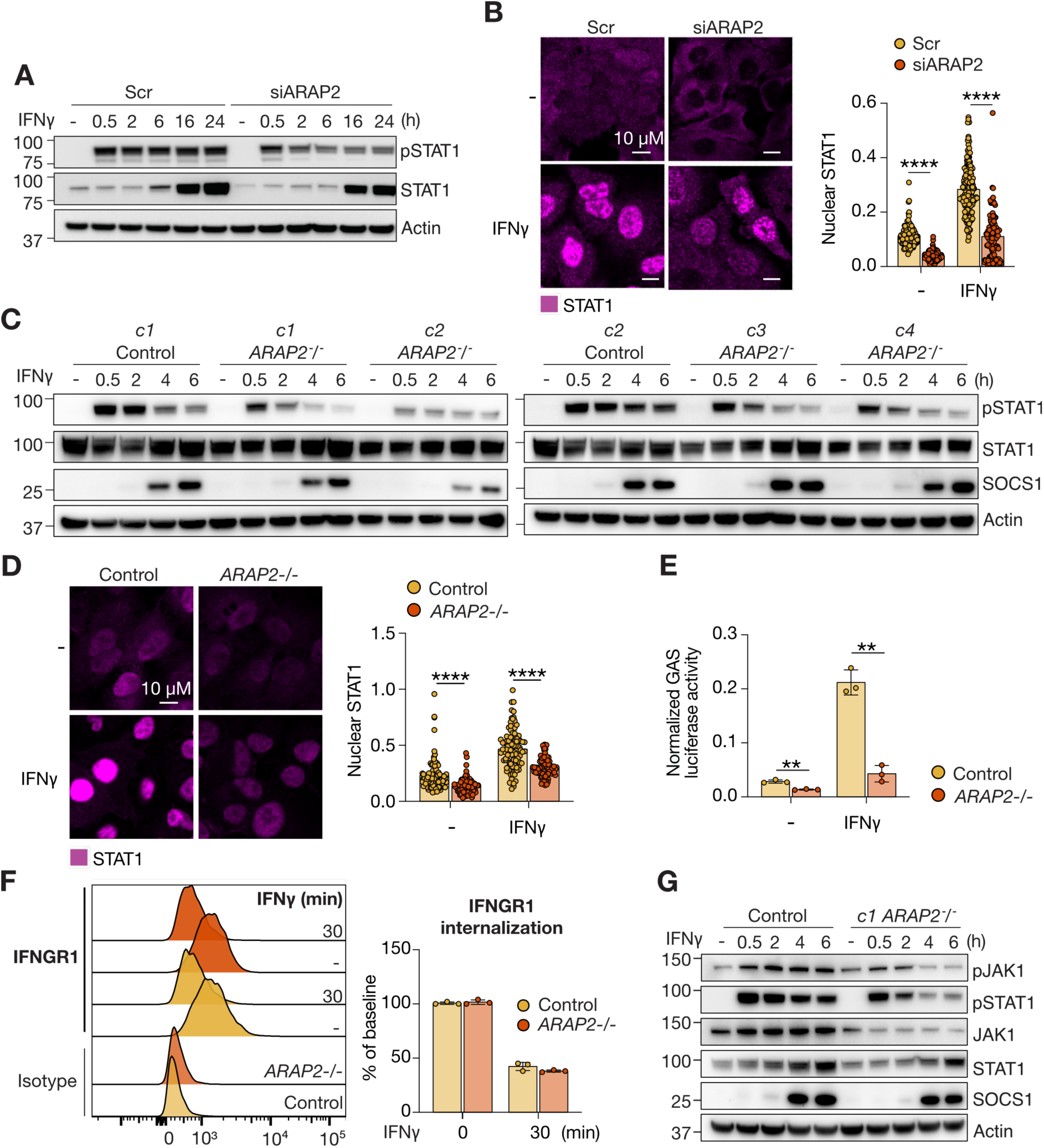
ARAP2 is required for effective IFNγ signaling. **A:** A549 cells were transfected with scramble or ARAP2-targeting siRNA. Cells were treated with IFNγ (100 ng/mL) for times indicated and lysates analyzed by SDS-PAGE and immunoblot. **B:** A549 cells were transfected with scramble or ARAP2-targeting siRNA. Cells were treated with 100 ng/mL IFNγ for 30 min prior to fixation and staining with STAT1 antibody, and analysis using a Zeiss LSM 880 microscope. Quantification of nuclear STAT1. n=50-70 cells. Mean -/+ S.D. Unpaired t-test. **C:** Control single guide (sg)RNA and *ARAP2^-/-^* A549 clonal cell lines were treated with IFNγ (100 ng/mL) for the times indicated and lysates analyzed by SDS-PAGE and immunoblot. **D:** Control sgRNA and *ARAP2^-/-^* A549 cells were treated with 100 ng/mL IFNγ for 30 min prior to fixation. Fixed cells were stained with STAT1 antibody and analyzed using a Zeiss LSM 880 microscope. Quantification of nuclear STAT1. n=50-70 cells. Mean -/+ S.D. Unpaired t-test. **E:** WT and *ARAP2^-/-^* HEK293T cells were co-transfected with GAS firefly reporter and renilla luciferase constructs in triplicate. 24 h post-transfection, cells were treated with IFNγ (100 ng/mL) for 16 h, and then analyzed for luminescence. Mean -/+ S.D. Unpaired t-test. **F:** Control sgRNA and *ARAP2^-/-^* A549 clones were treated with IFNγ (100 ng/mL) for the times indicated and stained with anti-IFNGR1 antibody conjugated to PE, prior to fixation and data collection using a Fortessa 1 Flow cytometer. **G:** Control sgRNA and *ARAP2^-/-^* A549 clonal lines were treated with IFNγ (100 ng/mL) for the times indicated and lysates analyzed by SDS-PAGE and immunoblot. **A-G:** Representative of three independent experiments. See Supplementary Figure 2 for repeat experiments and Sanger/Next Generation Sequencing analysis confirming CRISPR deletions.

Genetic deletion of *ARAP2* in A549 clonal cell lines (*ARAP2^-/-^*; confirmed by Next Generation Sequencing; **Supplementary** Figure 2E) also reduced the IFNγ-pSTAT1 response, compared to control clonal cell lines containing scrambled guides, validating the results with siRNA **(Figure 2C, Supplementary** Figure 2F). Similarly, deletion of *ARAP2* resulted in a significant decrease in STAT1 nuclear localization, and this was again evident before and after IFNγ treatment **(Figure 2D, Supplementary** Figure 2G **& H)**. The baseline changes in STAT1 nuclear localization may reflect low level activation of anti-viral pathways and autocrine production of IFN, in response to the exogenous siRNA and DNA introduced into the cell lines.

In addition, genetic deletion of *ARAP2* in HEK293T cells (confirmed by PCR and Sanger sequencing, **Supplementary** Figure 2I) also reduced IFNγ-induced pSTAT1 **(Supplementary** Figure 2J **& K)**. To assess STAT1 transcriptional activity, WT and *ARAP2*-deficient (*ARAP2^-/-^*) HEK293T cells were transiently transfected with an IFNγ activated site (GAS) reporter construct and stimulated with IFNγ for 16 h. GAS luciferase activity was reduced in *ARAP2^-/-^* HEK293T cells compared with control cells **(Figure 2E, Supplementary** Figure 2L **& M)**, indicating the reduced pSTAT1 and STAT1 nuclear accumulation observed in ARAP2-deficient cells was sufficient to impact STAT1 transcriptional activity.

Although ARAP2 was clearly required for maximal STAT1 signaling in response to IFNγ, it was unclear at which point ARAP2 impacted on the IFNγ signaling cascade. As ARAP2 was thought to be involved in endocytosis, we investigated whether loss of ARAP2 impacted IFNGR1 surface expression. However, no differences in IFNGR1 levels were observed in *ARAP2^-/-^* A549 cells, either at baseline or after 30 min IFNγ stimulation **(Figure 2F)**. Given that SOCS1 directly regulates JAK1 activity, we next examined whether loss of ARAP2 affected JAK1 phosphorylation. Interestingly, both phosphorylated and total JAK1 were reduced in *ARAP2^-/-^* cells **(Figure 2G, Supplementary** Figure 2N). *JAK1* is a known IFNγ-induced gene (INTERFEROME database^45^). It is possible that reduced baseline IFN signaling in the absence of ARAP2 resulted in reduced JAK1 levels. Alternatively, more SOCS1 may now be available to interact with JAK1 and in addition to inhibiting JAK1 kinase activity, may be enhancing ubiquitination of JAK1, directing it to proteasomal degradation.

Overall, these data suggested that ARAP2 regulated responses to IFNγ at the level of the JAK kinases, and that reduced JAK1 activity was responsible for a subsequent decrease in STAT1 phosphorylation, nuclear translocation and transcriptional activity.

SOCS1 also regulates type I and III IFN signaling^32, 42^, however, there were no differences in IFNα- or IFNƛ-induced pSTAT1 or pSTAT2 in *siARAP2* or *ARAP2^-/-^*A549 cells, compared with control cells **(Supplementary** Figure 3A-C). Additionally, there were no marked differences in IFNα-induced pSTAT1/pSTAT2 detection by immunoblotting or IFNα-induced-reporter activity in ARAP2-deficient HEK293T cells **(Supplementary** Figure 3D-E). Collectively, these data demonstrated that ARAP2 was a positive and selective modulator of type II IFNγ signaling.

### ARAP2 promotion of IFNγ signalling requires Tyr415 in ARAP2 and SOCS1

To investigate whether SOCS1 was required for the reduced IFNγ responses in the absence of ARAP2, ARAP2 was depleted from WT and *SOCS1^-/-^*A549 cells using siRNA. As in previous experiments, siRNA depletion of ARAP2 resulted in reduced pJAK1 and pSTAT1 following IFNγ treatment **(Figure 3A, Supplementary** Figure 4A **& B)**, however in contrast to *ARAP2* gene deletion, there were no differences in total JAK1. SOCS1 levels were elevated in *siARAP2* cells treated with IFNγ. Importantly, depletion of ARAP2 had no impact on pSTAT1 in *SOCS1^-/-^* A549 cells, suggesting that SOCS1 was required for ARAP2 to regulate IFNγ signaling. Consistent with this hypothesis, there was no difference in STAT1 nuclear accumulation following IFNγ treatment of *ARAP2*-depleted *SOCS1^-/-^* A549 cells **(Figure 3B, Supplementary** Figure 4C **& D)**. However, the decreased STAT1 nuclear localization observed at baseline also occurred in *SOCS1^-/-^ siARAP2* cells, albeit with some variation between individual experiments. These data suggested that the reduced IFNγ response observed in *ARAP2^-/-^* cells was largely mediated by SOCS1.

**Figure 3:**
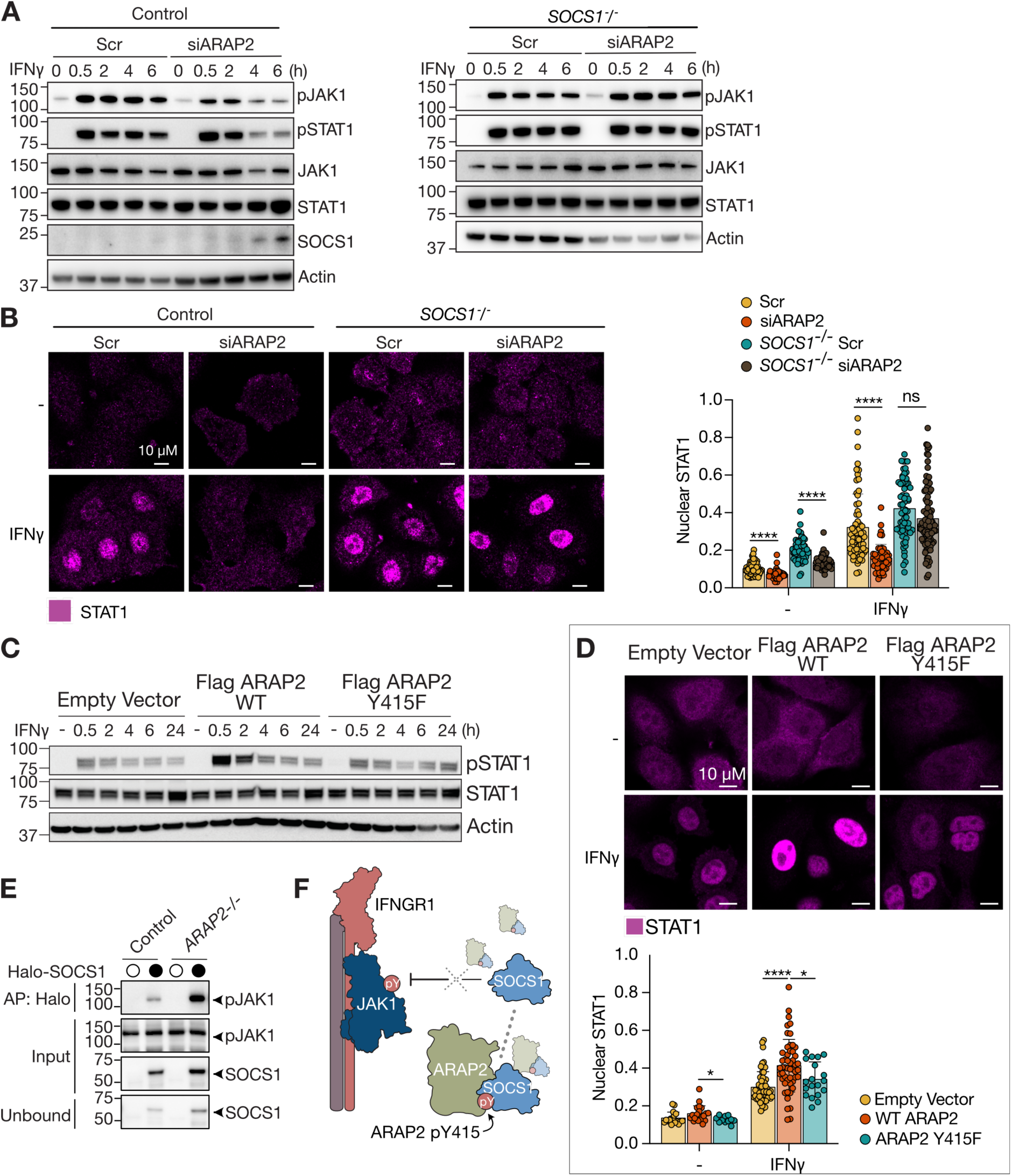
ARAP2 restricts the ability of SOCS1 to inhibit signaling. **A:** WT and *SOCS1^-/-^* A549 cells were transfected with scramble or ARAP2-targeting siRNAs and treated with IFNγ (100 ng/mL) for the times indicated. Lysates were analyzed by SDS-PAGE and immunoblot. **B:** WT and *SOCS1^-/-^* A549 cells were transfected with scramble or ARAP2-targeting siRNAs and treated with IFNγ (100 ng/mL) for 30 min prior to fixation. Fixed cells were stained with anti-STAT1 antibody (magenta) and DAPI (not shown) and analyzed using a Zeiss LSM 880 microscope. Quantification of nuclear STAT1. n=50-70 cells. Mean -/+ S.D. Unpaired t-test. **C:** A549 cells were transiently transfected with empty vector (EV), Flag-ARAP2 WT or Flag-ARAP2 Y415F constructs. 48 h post-transfection, cells were divided for an IFNγ time course and stimulated with IFNγ (100 ng/mL) for times indicated and analyzed by SDS-PAGE and immunoblot. **D:** A549 cells were transiently transfected with Empty vector (EV), Flag-ARAP2 WT and Flag-ARAP2 Y415F constructs and stimulated with IFNγ (100 ng/mL) for 30 min prior to fixation and incubation with anti-STAT1 antibody (magenta) and DAPI (not shown) for confocal analysis using the Zeiss LSM 880 microscope. Quantification of nuclear STAT1. n=50-70 cells. Mean -/+ S.D. Unpaired t-test. **E:** Control sgRNA and *ARAP2^-/-^* A549 cells stably expressing a doxycycline inducible construct of Halo-SOCS1 were treated with doxycycline overnight and pervanadate (100 µM) for 30 min prior to lysis. Halo-Flag-SOCS1 was enriched with HaloLink resin and the presence of endogenous pJAK1 and ARAP2 analyzed by SDS-PAGE and immunoblot (left panel). Note that Halo-SOCS1 is covalently linked to the beads via the Halo ligand. The input and unbound fractions confirm enrichment of SOCS1 during affinity purification (AP). **F:** Schematic depicting hypothesis that ARAP2 promotes IFNγ signaling by restricting SOCS1 from inhibiting pJAK1. **A-E:** Representative of three independent experiments. See Supplementary Figures 4-5 for repeat experiments.

We next investigated whether mutation of the key SOCS1-interacting tyrosine in ARAP2 to phenylalanine (ARAP2-Y415F) impacted the IFNγ response. A549 cells were transiently transfected with empty vector (EV), Flag-ARAP2-WT or Flag-ARAP2-Y415F constructs and stimulated with IFNγ. Exogenous expression of Flag-ARAP2-WT, but not Flag-ARAP2-Y415F, enhanced IFNγ-induced STAT1 phosphorylation **(Figure 3C, Supplementary** Figure 5A-C). Furthermore, exogenous expression of Flag-ARAP2-WT, but not Y415F, resulted in a two-fold increase in STAT1 nuclear localization following IFNγ treatment **(Figure 3D, Supplementary** Figure 5D **& E)**. These data suggested that Tyr415 in ARAP2 was also required to promote IFNγ signaling.

We hypothesized that ARAP2 restricted the ability of SOCS1 to inhibit JAK1. If this was correct, we predicted that the SOCS1:JAK1 interaction would be enhanced in the absence of ARAP2. A doxycycline inducible construct of Halo-SOCS1 was stably introduced into scramble and *ARAP2^-/-^* A549 cells. Cells were treated with pervanadate to maximize tyrosine phosphorylation prior to affinity purification of Halo-SOCS1 using HaloLink resin. Immunoblot analysis showed increased expression of Halo-SOCS1 protein in *ARAP2^-/-^*cells, with a corresponding increase in pJAK1 enrichment **(Figure 3E, Supplementary** Figure 5F **& G)**. The increased Halo-SOCS1 appeared related to pervanadate treatment, given that SOCS1 levels were not consistently elevated in *ARAP2* deleted clones **(Figure 2C)**. Greater SH2 interaction with a phosphorylated target is likely to stabilize SOCS1 protein, and although the results are consistent with ARAP2 sequestering SOCS1 away from pJAK1 **(Figure 3F)**, this data remained inconclusive. Despite this caveat, the data demonstrated that SOCS1 and ARAP2-Y415 were required for ARAP2 to potentiate IFNγ signaling.

### Mutation of ARAP2-Y413F attenuates disease severity in response to influenza infection

Influenza infection in mice results in cytokine-mediated weight loss followed by an adaptive immune response and viral clearance^46^. Although IFNγ is primarily a macrophage-activating factor, as unequivocally shown in humans genetically lacking IFNγ, it does exhibit anti-viral activity under certain experimental conditions in mice^1^. It can also drive detrimental inflammatory pathology in viral infection, with various studies showing that loss of IFNγ in mice protects against weight loss during influenza infection^47, 48, 49^.

To investigate the role of IFNγ and ARAP2-Y413 in influenza infection, we generated C57BL/6J (WT) mice that either lacked ARAP2 (*Arap2^-/-^*) or carried a germ-line mutation of Y413F (*Arap2^Y413F/Y413F^*; equivalent to human ARAP2-Y415). Intercrossing heterozygous mice produced offspring in mendelian ratios, and homozygous mutant mice were fertile and displayed normal size and weight, with no overt phenotype. Analysis of thymus, spleen, lymph node and bone marrow, did not reveal any major perturbations in T cell development or mature immune subsets under homeostatic conditions **(Supplementary** Figure 6A-C).

We next investigated the cell-types in the lung that expressed SOCS1 during influenza A virus (A/Puerto Rico/8/1934 (H1N1); PR8) infection. Mice carrying a germline insertion of a *Halo-Socs1* fusion sequence under regulatory control of the endogenous *Socs1* promoter (*Halo-Socs1^KI/KI^*)^50^, were utilized to examine expression of Halo-SOCS1 in the lung following PR8 viral infection. Lungs were harvested from WT and *Halo-Socs1^KI/KI^*mice infected with PR8 virus at day 7 post-infection, and TMR-staining performed in lung homogenates processed to single cell suspensions. Day 7 was selected based on our previous studies and those of others showing maximum weight loss at this timepoint^49^. TMR staining revealed expression of Halo-SOCS1 protein in most cell populations examined, except for neutrophils (**Figure 4A, Supplementary** Figure 6D). Halo-SOCS1 expression relative to background staining in WT cells was highest in NK cells, CD4^+^ and CD8^+^ T cells, and B cells. In general, Halo-SOCS1 protein expression was consistent with *Socs1* transcript levels in immune subsets in the lungs of C57BL/6 mice at day 7 post-PR8 infection (single cell RNA sequencing data^49^), with high *Socs1* levels in lung-infiltrating neutrophils, B cells, CD4^+^ and CD8^+^ T cells, and to a lesser extent, NK cells **(Figure 4B-D)**. Expression of *Arap2* transcript was also highest in NK cells, T cells and B cells, with very little expression in neutrophils and monocytes **(Figure 4B-D)**. Overlapping expression of *Socs1* and *Arap2* in NK, T and B cells, suggested there might be a functional role for their interaction in these cells during influenza infection.

**Figure 4:**
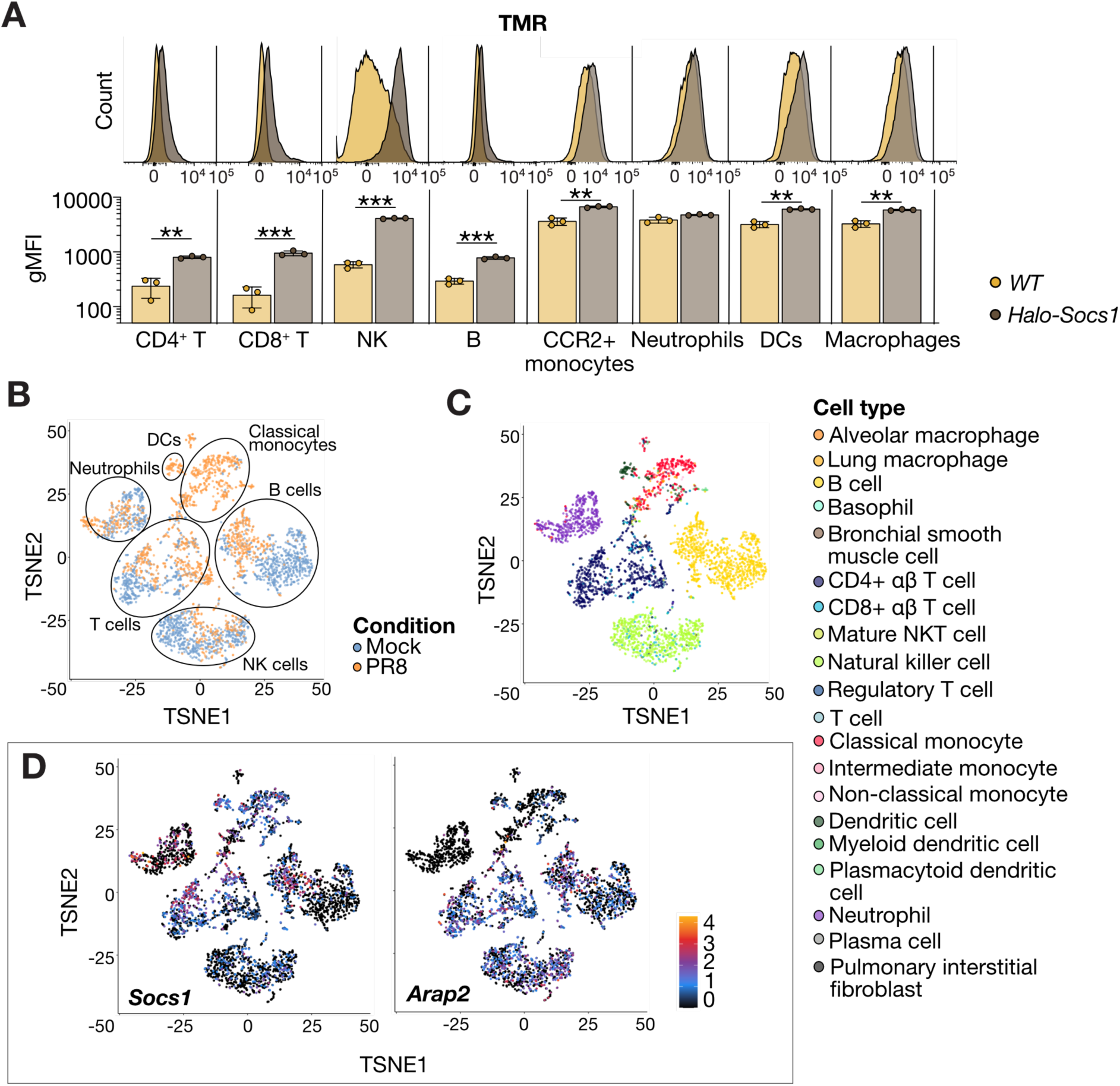
Socs1 and *Arap2* are expressed in PR8-infected immune cells. **A:** SOCS1 expression in immune cell subtypes. Wild-type (WT) and *Halo-Socs1^KI/KI^* male mice were infected i.n. with 40 PFU influenza virus H1N1 PR8 and lungs harvested at day 7 post-infection. Lungs were digested and viable, single cells analyzed for expression of Halo-SOCS1 protein by TMR-ligand and flow cytometry. Upper panels show representative histograms. Lower panels show gMFI from 3 different mice. Mean ± S.D. \**p*< 0.05; \*\**p*<0.01; \*\*\**p*<0.001, using an unpaired *t*-test. Example gating strategy is shown in Supplementary Figure 6D. **B-D:** Analysis of single cell RNA seq data from lungs of C57BL/6J mice at day 7 post-infection with PR8 virus. Data were re-analyzed from Schmit *et al*. 2022^49^. TSNE plots showing: **(B)** Various immune cell populations and highlighting the influx of cells in response to PR8 infection (orange); **(C)** Identified cell types; **D)** *Socs1* and *Arap2* expression.

WT, *Ifng^-/-^, Arap2^-/-^* and *Arap2^Y413F/Y413F^*C57BL/6J mice were infected with PR8 influenza virus and weight loss monitored daily for 10 days. In-line with previous studies, *Ifng^-/-^* mice lost significantly less body weight than their WT counterparts following influenza infection, and this was evident from day 6 post-infection **(Figure 5A, Supplementary** Figure 6E), confirming a pathogenic role for IFNγ during influenza infection in C57BL/6J mice. *Arap2^-/-^* and *Arap2^Y413F/Y413F^* mice also displayed less weight loss compared to WT mice following PR8 infection **(Figure 5B & C, Supplementary** Figure 6F **& G)**, consistent with reduced IFNγ production or signaling.

**Figure 5:**
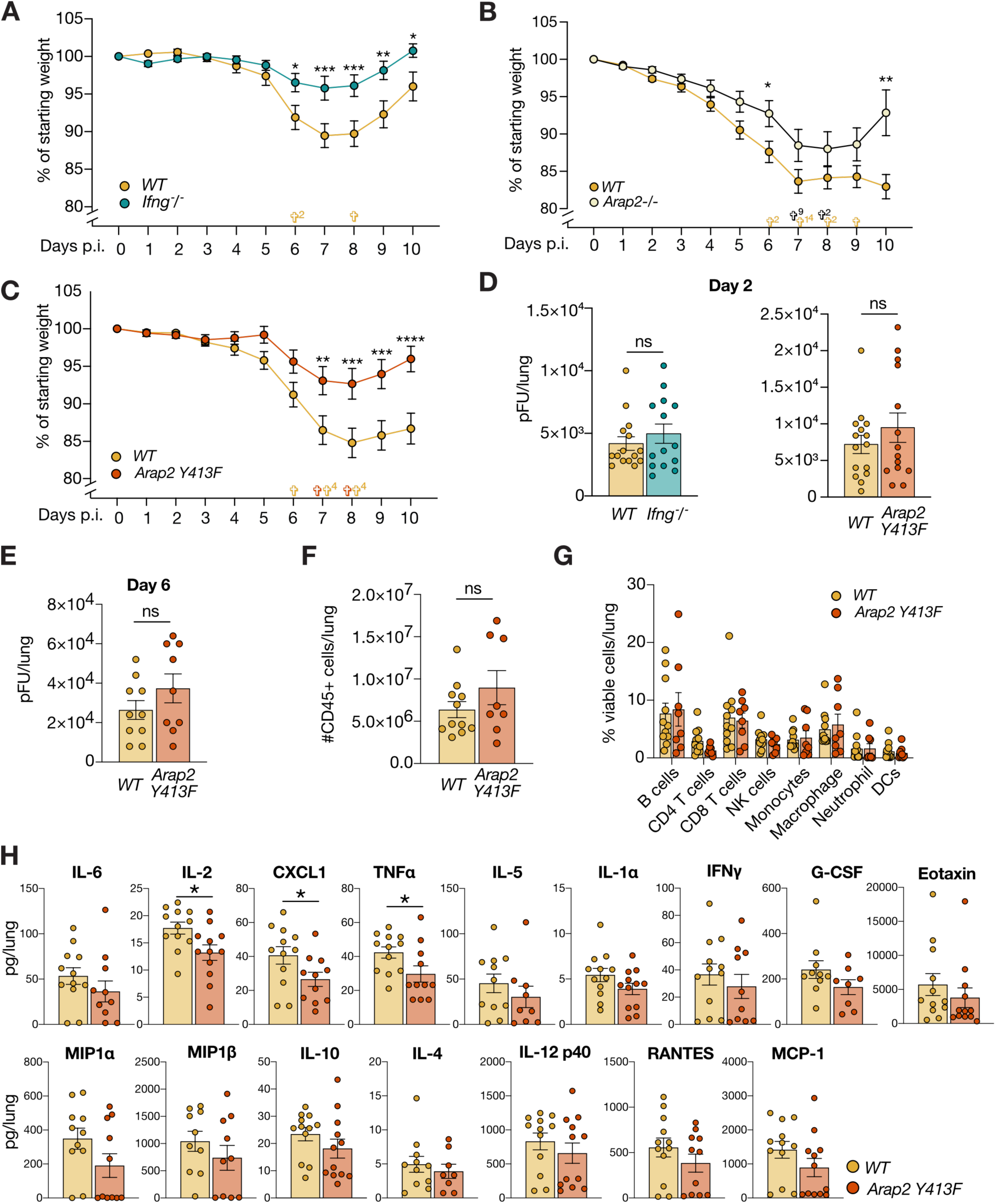
Arap2Y413F mice are protected from PR8 influenza virus-induced weight-loss. **A-C:** Wild-type (WT) and **(A)** *Ifng^-/-^*, **(B)** *Arap2^-/-^* and **(C)** *Arap2^Y413F^* male mice were infected i.n. with 40 PFU influenza virus H1N1 PR8 and weight loss monitored for 10 days. Uindicates mice that lost more than 20% of their initial body weight and were euthanized. Superscript indicates number of mice. The last recorded weight is included in subsequent days to normalize data. Data are combined from two independent experiments. n = 18-20. Mean ± S.E.M. \**p*< 0.05; \*\**p*<0.01; \*\*\**p*<0.001 as analyzed by two-way ANOVA. **D-E:** Viral load (PFU/lung) in lung homogenates at day 2 **(D)** and day 6 **(E)** post-infection was determined in an independent experiment by standard plaque assay (right panel). Mean ± S.E.M. ns=not significant. **F:** Absolute numbers of CD45+ immune cells per lung at day 7 post-infection analyzed by flow cytometry. Data are combined from two independent experiments. WT: n = 11; *Arap2^Y415F^*: n = 9. Mean ± S.E.M. **G:** Composition of immune cell subsets in the lung at day 7 post-infection analyzed by flow cytometry. Data are combined from two independent experiments. WT: n = 11; *Arap2^Y415F^*: n = 8. Mean ± S.E.M. **F and G:** Example gating strategy in Supplementary Figure 6D. **H:** Cytokine and chemokine levels in lung homogenates at day 6 post-infection were assessed using a BioPlex assay. Combined data from two independent experiments. n = 8-12. Mean ± S.E.M. *p<0.05 as analyzed by two-way ANOVA.

Further analysis was focused on *Arap2^Y413F/Y413F^* mice. Viral loads in the lungs of *Ifng^-/-^* and *Arap2^Y413F/Y413F^* mice were comparable to WT mice at day 2 post-infection **(Figure 5D)**, suggesting that the reduced weight loss did not result from reduced viral entry and/or initial viral replication. Viral loads at day 6 post-infection tended to be higher in *Arap2^Y413F/Y413F^* mice but were not significantly different from WT mice **(Figure 5E).** We next examined immune cell composition in the lungs of WT and *Arap2^Y413F/Y413F^*mice at day 7 post influenza infection, but did not observe any differences in numbers of CD45+ immune cells or skewing in the proportion of immune cell subsets **(Figure 5F & G, Supplementary** Figure 6D). However, levels of pro-inflammatory cytokines and chemokines were generally reduced in lungs of *Arap2^Y413F/Y413F^* mice compared to WT mice at day six post-infection. In particular, IL-2, TNFa and CXCL1 were significantly reduced in lungs from *Arap2^Y413F/Y413F^* mice **(Figure 5H)**. These data suggested that the reduced weight loss observed in *Arap2^Y413F/Y413F^* mice resulted from a dampened cytokine storm in the lungs, which was not related to reduced viral load or immune cell infiltration.

These data revealed that ARAP2 and Tyr413 in ARAP2 had an important role in regulating the inflammatory response to influenza virus infection *in vivo*. Given the complex interplay between cytokines, chemokines and immune cells in the lung during influenza infection, it is difficult to prove a direct link between ARAP2 and IFNγ during the control of influenza infection in mice. However, taken together our *in vitro* and *in vivo* data suggest that loss of ARAP2 or mutation of ARAP2-Y413, results in reduced responses to IFNγ, consequentially leading to a reduced inflammatory response and protection against influenza-induced weight loss.

## DISCUSSION

Here, we discovered ARAP2 to be a novel regulator of responses to IFNγ that fine-tunes SOCS1-mediated inhibition of JAK1. Tyr415 in human ARAP2 was identified as a high-affinity binding site for the SOCS1-SH2 domain. Depletion of ARAP2 using either siRNA or CRISPR-Cas9 gene editing dampened cellular responses to IFNγ, which was rescued by deletion of the *SOCS1* gene. Furthermore, exogenous expression of ARAP2, but not ARAP2-Y415F augmented the response to IFNγ, resulting in heightened STAT1 tyrosine-phosphorylation, nuclear accumulation and transcriptional activity. Finally, mice carrying a germline deletion of *Arap2* or mutation of the analogous Y413 in *Arap2* (Y413F), displayed reduced inflammation and disease severity in response to influenza infection, which was comparable to that observed in *Ifng^-/-^* mice. We propose that during IFNγ signaling, ARAP2-Y415 restricts the ability of SOCS1 to inhibit JAK1.

ARAP2 is a 193 kDa member of the ADP-ribosylation factor (Arf) GTPase-activating protein (GAP) family (which includes ARAP1 and ARAP3) and was discovered based on homology with other ArfGAPs^51^. ARAP2 is comprised of a sterile alpha-motif (SAM), an Arf-GAP domain, an ankyrin repeat, an inactive Rho-GAP domain, a Ras-association domain, and five pleckstrin homology (PH) domains. Very few publications have investigated ARAP2 structure-function, however a basic understanding can be inferred from its domain architecture. The N-terminal SAM domain facilitates binding of phosphatidylinositol lipids to the PH domains, which in turn localize ARAP2 to the plasma and endosomal membranes, and focal adhesions^51,52^. The ARAP2-Arf-GAP domain hydrolyses GTP to GDP, with high selectivity for Arf6^52^, a GTP binding protein that is a central component of the regulatory machinery controlling endocytic trafficking^53^, colocalizing with Arf6 in the early endosome^54^. Tyrosine 415 is located in a predicted unstructured region on one face of the structured domains (this paper), suggesting it is accessible for both phosphorylation and SH2 binding. The role of ARAP2 in the regulation of immunity or IFNγ signaling had not been investigated prior to this study.

IFNγ signaling has multiple tiers of spatial and temporal regulation that act in concert to maintain the appropriate level of signaling. One such mechanism is endocytic trafficking, whereby internalization is required for robust IFNγ signaling activation. We initially hypothesized that ARAP2 regulated the response to IFNγ through internalization of the IFNGR complex and/or endocytic trafficking; but did not observe any differences in IFNGR expression on the surface of cells lacking ARAP2. Furthermore, there were no significant differences in influenza viral load in the lungs of infected ARAP2-Y413F mice at day 2 or day 6 p.i., evidence that the reduced weight loss in *Arap2^Y413F^* mice was not due to differences in initial viral uptake, trafficking or replication.

IFNγ functions principally as a MAF and exerts only modest antiviral activity^1^. In fact, IFNγ has a pathogenic role in mice following influenza infection, with IFNγ-deficient mice displaying reduced disease severity, manifested as reduced weight loss (this paper)^49^. Type I and III IFNs are induced in the early stages of influenza infection and are required for effective viral immunity^55^, whereas type II IFNγ levels peak at day 7 p.i.^49^. We (this paper) and others^49^ have shown that SOCS1 is present in multiple immune cell types during the period of maximal IFNγ and the associated weight loss (day 7 p.i.), consistent with SOCS1 protecting against IFNγ-mediated inflammation. The apparently restricted role of ARAP2 to type II IFN signaling is supported by our *in vitro* studies showing that ARAP2 was required for effective IFNγ, but type I or III IFN, signaling responses. Thus, our discovery that both *Arap2^-/-^* and *Arap2^Y413F^* mice also displayed reduced weight loss and production of inflammatory cytokines during influenza infection, supported the idea that ARAP2-deficiency or mutation increased SOCS1 activity and protected against IFNγ-driven inflammation.

Interestingly, the ARAP2 Y415F mutation phenocopied full *ARAP2* deletion *in vitro*, resulting in reduced responses to IFNγ. This was surprising, given the various domains present in ARAP2 with presumably independent biological functions, and indicates Y415 is critical for the functional interplay between ARAP2 and IFNγ signaling responses. While our data support ARAP2-Y413/5 sequestering SOCS1 to limit its inhibition of JAK1, it remains possible that SOCS1 regulates ARAP2 independently of its interaction with JAK, and this indirectly impacts responses to IFNγ. However, our experiments have not provided any evidence to suggest that SOCS1 regulates ARAP2 levels. Indeed, the SOCS1-SOCS box only weakly interacts with Cullin-5, the scaffold protein involved in SOCS box-mediated ubiquitination^56^.

Given that SOCS1 negatively regulates signaling downstream of type I^35^, II^16^ and III^36^ IFN, what determines the apparent selective requirement for ARAP2 in IFNγ signaling remains an elusive question, although this may be linked to ARAP2 expression, or the cellular context for phosphorylation (and de-phosphorylation) of ARAP2 Y413/5. Our data suggested ARAP2 was phosphorylated in the absence of IFNγ stimulation (evidenced by SOCS1:ARAP2 interaction without cytokine activation). It is possible that the SOCS1:ARAP2 interaction is temporally controlled by dephosphorylation of ARAP2, and this is regulated by IFNγ signaling. Future studies identifying the kinases and phosphatases that regulate ARAP2 Y415 phosphorylation would provide critical mechanistic insight.

There are relatively few examples of proteins that directly regulate the SOCS proteins, other than by inducing their transcript. SOCS3 interaction with the caveolae protein, cavin-1, localizes SOCS3 to the plasma membrane and is essential for SOCS3 function^57^. Loss of cavin-1 enhanced STAT3 phosphorylation and abolished SOCS3 inhibition of IL-6 signaling. Moreover, Linossi et al.^58^, discovered an exosite on the SOCS2-SH2 domain which, when bound to a non-phosphorylated peptide (F3), enhanced SH2 affinity for canonical phosphorylated ligands. These additional protein-protein interactions, now including ARAP2, may be an under-appreciated facet of SOCS protein activity.

In this study, we discovered that ARAP2 was critical for robust responses to IFNγ, strictly and selectively regulating SOCS1 activity via an interaction between the SOCS1-SH2 domain and ARAP2-Y415. Moreover, we uncovered a role for ARAP2 in control of IFNγ-driven inflammation. Traditionally, steroidal and non-steroidal anti-inflammatory drugs have been used to treat severe infections such as Influenza, however these are often not effective or are associated with severe side-effects^52^. More recently, immunomodulators such as anti-IL-6 receptor antibodies^53^ and JAK inhibitors^5459^, have shown promise for the treatment of inflammatory viral diseases such as COVID-19^60^. The data presented in this study suggest that immunomodulation of IFNγ-driven inflammation, such as through SOCS1 agonists or by disruption of the ARAP2:SOCS1 interaction, may be useful for treatment of inflammatory diseases such as severe Influenza A.

## ACKNOWLEDGEMENTS

We acknowledge the Wurundjeri people of the Kulin nation as the traditional owners and guardians of the land on which the majority of the work was performed. The authors thank Prof. Paul Hertzog (Monash University) for helpful discussions, and Natasha Blasch, Sophia Russo, Natasha Chow, Anja Lu, Jessica Martin, Keti Florides, Eren Loza and Natalia MoraTorres for mouse husbandry. N.K. was supported in part by an Australian government Research Training Program Scholarship and a Fulbright Scholarship. J.J.B was supported by an Australian Government National Health and Medical Research Council (NHMRC) Research Fellowship (1121755). K.L. was supported by a Melbourne Research Scholarship (University of Melbourne), G.M.B. by an Australian government Research Training Program Scholarship, and L.G.M.G. by a Walter and Eliza Hall Institute International PhD Scholarship. This work was supported in part through NHMRC Ideas Grant 2011761, Victorian State Government Operational Infrastructure Support and the Australian Government NHMRC Independent Research Institutes Infrastructure Support Scheme (IRIISS). The generation of *Arap2^Y413F/Y413F^* mice used in this study was supported by Phenomics Australia and the Australian Government through the National Collaborative Research Infrastructure Strategy (NCRIS) program.

## AUTHOR CONTRIBUTIONS

**NK:** Conceptualized and led the study. Designed, executed, and analyzed experiments. NK wrote the original draft and co-reviewed and edited the paper with SEN.

**KD:** Led the generation of different ARAP2-Y413F and ARAP2-null mice founder lines and backcrossing, maintained mouse colonies, and produced ethics guidelines with SEN and LGMG. Performed the immunophenotyping of ARAP2-Y413F mouse strain, and analysis of immune cell populations during PR8 infection. Contributed to experiments that were not used in the final draft of the manuscript.

**GMB:** Helped generate and characterize A549 *SOCS1^-/-^*cells. Performed Halo-SOCS1 flow cytometry experiments and assisted with immunophenotyping.

**LGMG:** Processed organs for immunophenotyping and performed FACS analysis. Also produced ethics guidelines with KD and SEN.

**LFD:** Supervised experimental design and sample processing for mass spectrometry, acquired mass spectrometry data and performed peptide searching and data analysis.

**KL:** Generated protein for SPR experiments. Performed all competitive SPR assays. **AG:** Performed Bioplex cytokine assays and additional experiments that were not included in the final version of the manuscript.

**CA, BEW, BJB and BCD:** Performed intranasal *in vivo* PR8 influenza injections.

**CH:** Helped generate and characterize A549 *SOCS1^-/-^* cells and established the imaging pipeline, including quantification, for STAT1 microscopy.

**EL:** Performed genotyping of mice, and next generation sequencing for CRISPR-deleted cell lines.

**RM:** Provided critical reagents in the early phases of the study for experiments that informed but were not included in the final manuscript.

**AK:** Designed and supervised CRISPR/Cas9 gene editing strategy for mutation of ARAP2-Y413F in mice.

**ALG:** Performed bioinformatic analysis of Schmit et al., 2022 for *Socs1* and *Arap2* expression.

**JLC:** Provided reagents and advice on research direction. Supervised NK while performing ARAP2-deficient HEK293T immunoblot and luciferase assay experiments. Acquired project funding.

**SBD:** Designed the ARAP2 guides used to generate ARAP2-deficient HEK293T cells. Supervised the ARAP2 deletion in HEK293T and advised on IFN-response immunoblot and luciferase assay experiments. Provided cells and reagents. Acquired project funding.

**JJB:** Supervised SPR and isothermal titration calorimetry (ITC) experiments (ITC not included in final manuscript). Provided expression constructs and recombinant proteins.

**EML:** Generated recombinant protein. Supervised protein production and initial mass spectrometry experiment, which discovered the interaction between CIS and ARAP2.

**MDT:** Performed viral load plaque assays on lung homogenates derived from PR8 influenza infected mice.

**JRG:** Provided advice and supervised the *in vivo* PR8 influenza infections.

**SEN:** Conceived and supervised the study. Edited the manuscript. Acquired project funding.

All authors reviewed the manuscript.

## KEY RESOURCES TABLE

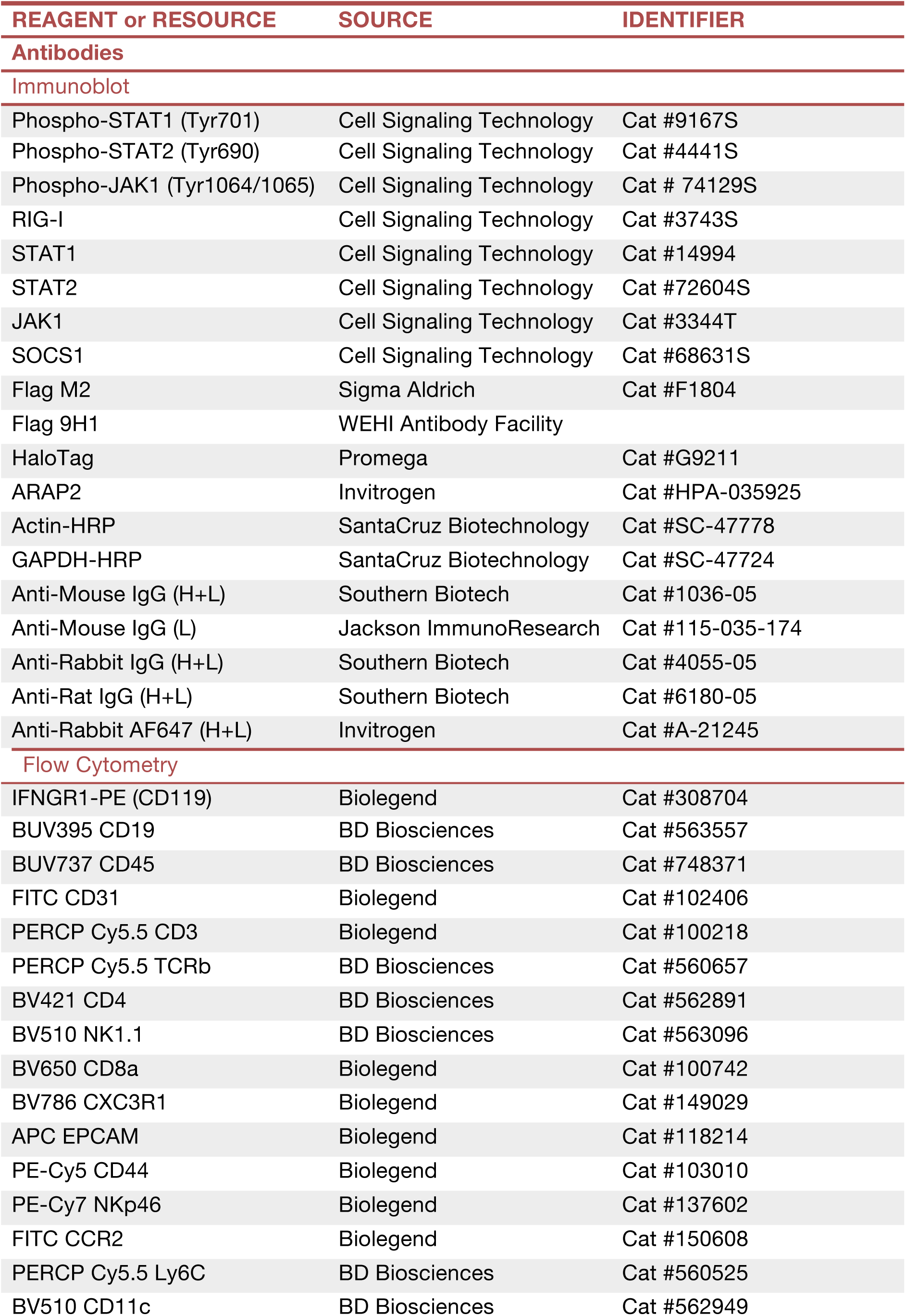
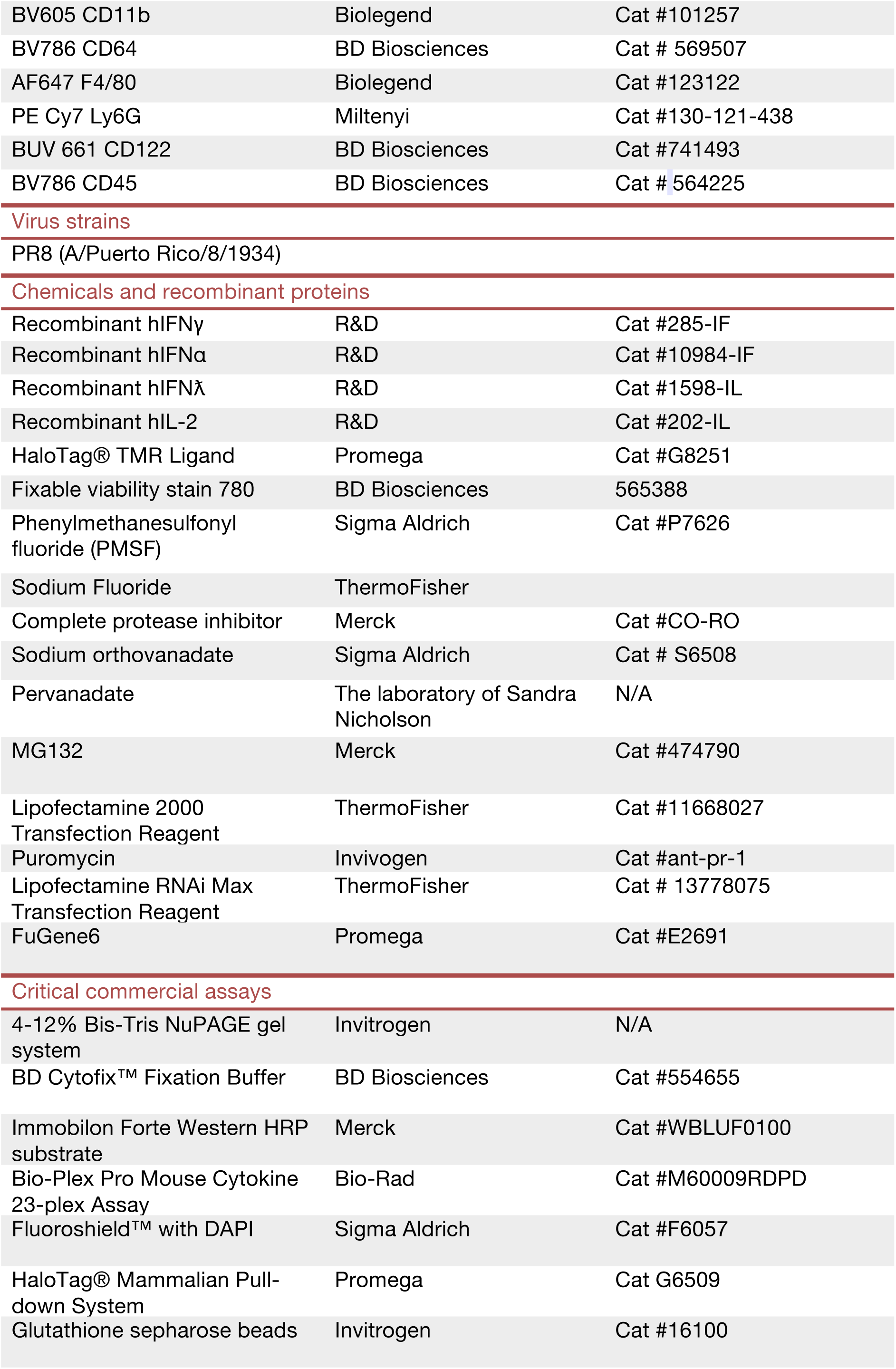
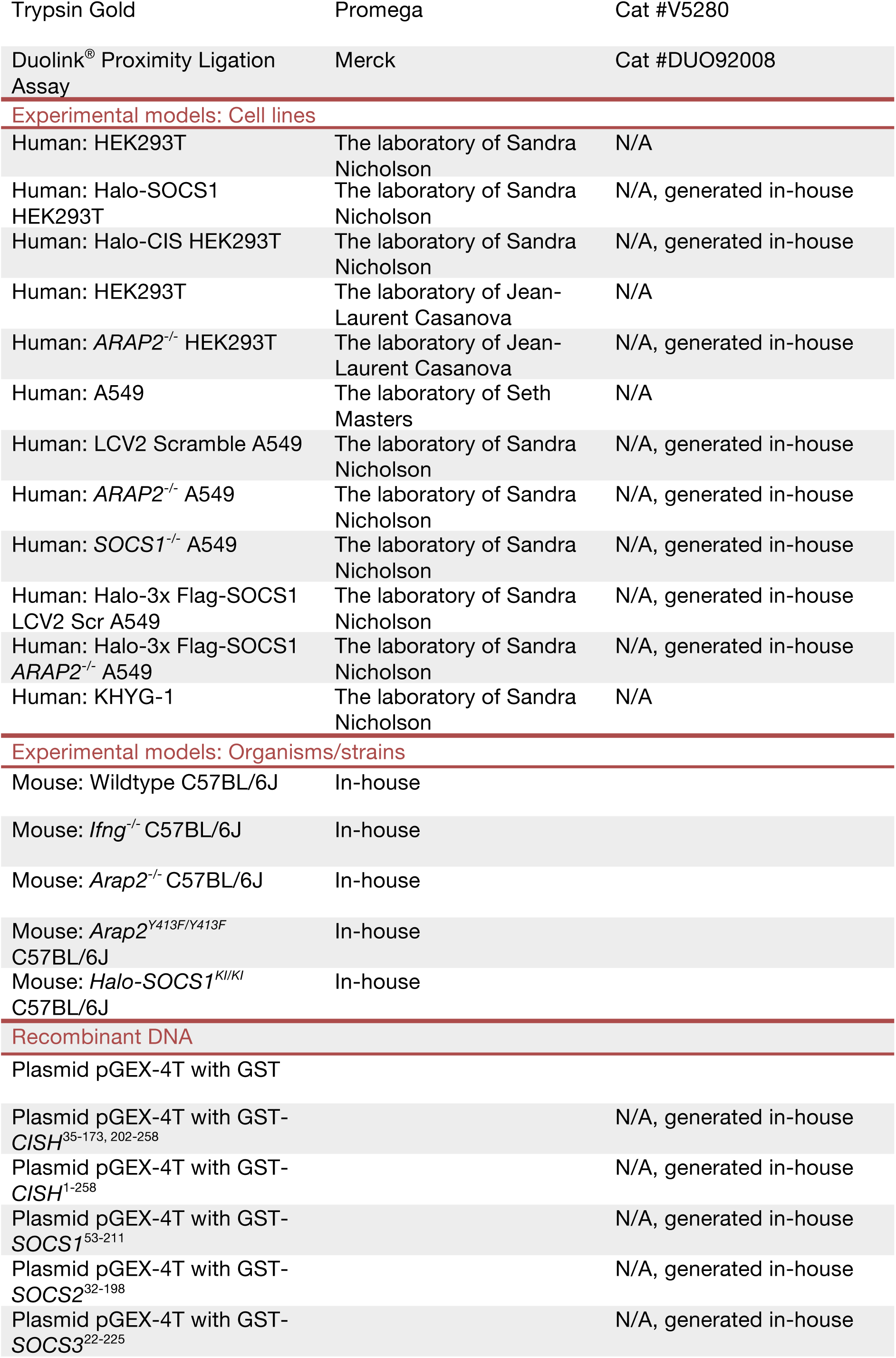
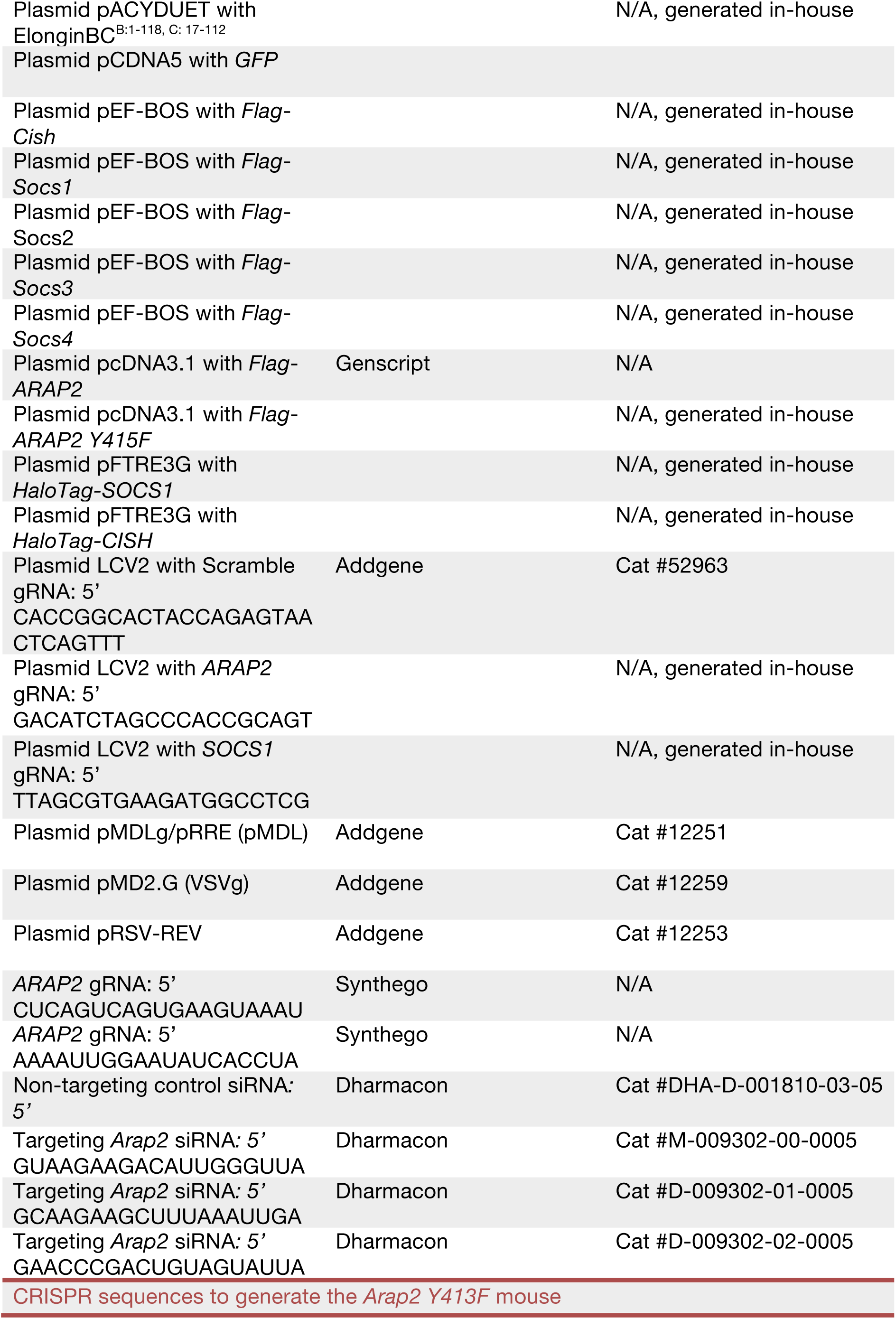
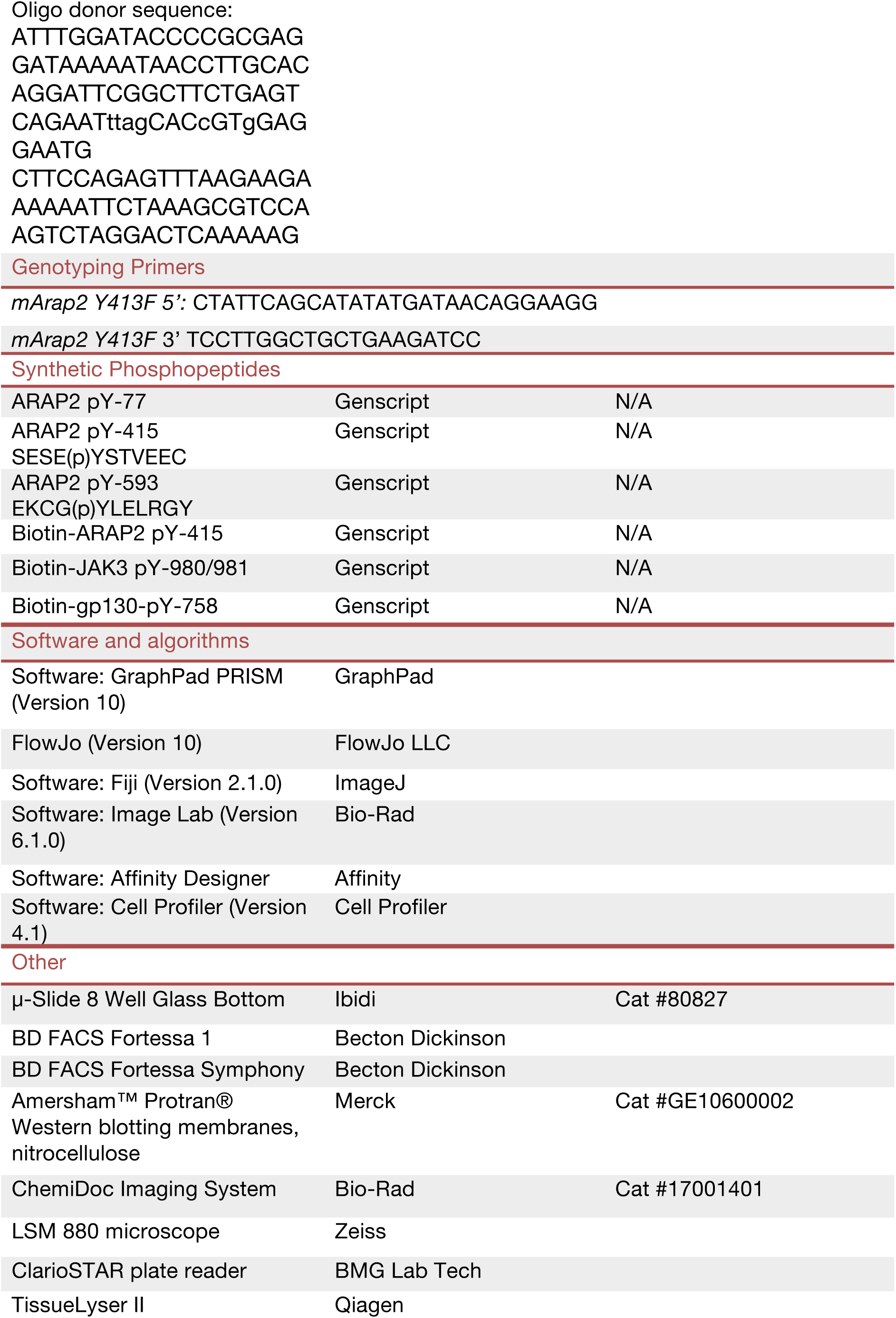

## RESOURCE AVAILABILITY

### Lead Contact

Further information and requests for resources and reagents should be directed to and will be fulfilled by the lead contact, Sandra E. Nicholson.

### Materials availability

All unique reagents generated in this study are available from the lead contact with a completed materials transfer agreement, with some exceptions due to commercial obligations.

### Data and code availability

- This paper does not report original code.
- Any additional information required to reanalyze the data reported in this paper is available from the lead contact upon request.

## EXPERIMENTAL MODEL AND SUBJECT DETAILS

### Mice

Mice were maintained on a C57BL/6J background under specific-pathogen-free conditions at the Walter and Eliza Hall Institute of Medical Research (WEHI), Australia.

Mice carrying a germline mutation of Tyr413 to Phe (Arap2-Y413F) in ARAP2 were generated on a C57BL/6J background by the MAGEC laboratory at the Walter & Eliza Hall Institute, as previously described [37]. 20 ng/µL Cas9 mRNA, 10 ng/µL single guide RNAs, and 40 ng/µL donor template were injected into fertilized one-cell stage embryos. The oligo donor of the sequence is provided in **KEY RESOURCES TABLE.** Founder mice were analyzed by next-generation sequencing (NGS) to confirm the correct sequence change. Mice carrying the mutation were backcrossed to C57BL/6J mice for three generations to eliminate potential off-target events, and next-generation sequencing repeated. Heterozygous mice were intercrossed to generate homozygotes, and litters were tattooed and sampled for genotyping at birth. Mice were genotyped by NGS using genomic DNA extracted from ear biopsies with the Direct PCR Lysis tail reagent (Viagen) and 5 mg/mL proteinase K (Worthington), according to the manufacturer’s instructions. Genotyping primers are shown in **KEY RESOURCES TABLE.** *Ifng^-/-^*^61^ *and Halo-Socs1 (Socs1^KI/KI^)* (Bidgood et al., 2024) mice have been described previously.

Experiments were performed at the Walter and Eliza Hall Institute (WEHI) in accordance with the NHMRC Australian code for the care and use of animals for scientific purposes. All experiments were approved by the WEHI Animal Ethics Committee (AEC 2021.002).

**PCR conditions:**

| Cycles | Temperature | Time |
| --- | --- | --- |
| 1 | 95°C | 2 min |
| 35 | 95°C | 30 s |
|  | 60°C | 30 s |
|  | 72°C | 30 s |
| 1 | 72°C | 5 min |

## METHOD DETAILS

### Expression vectors

All expression constructs are listed in the **KEY RESOURCES TABLE**.

Recombinant glutathione S-transferase (GST)-tagged SOCS proteins were co-expressed with Elongins B and C in *E. coli*. Production as a trimeric complex stabilizes the SOCS proteins. Expression and purification steps were performed as described^62^. For surface plasmon resonance assays, the GST-tag was removed by tobacco etch virus (TEV)-cleavage. Constructs encoding Flag-tagged SOCS proteins in a pEFBOS expression vector have been previously reported^63^. Constructs encoding hSOCS1 and hCIS with an N-terminal Halo tag were generated by polymerase chain reaction (PCR) to give fragments with BamHI and NheI restriction sites at the N- and C-termini respectively, and sub-cloned into the lentiviral Tet-On pfTRE3G puromycin expression vector. pcDNA3.1-Flag-hARAP2 (C-terminal DYKDDDDK) was purchased from Genscript. The ARAP2 Y415F mutant was generated using the Quikchange Lightening mutagenesis kit (Agilent Technologies), according to the manufacturer’s instructions. CRISPR oligonucleotides targeting ARAP2 were cloned into the LentiCRISPR-V2 vector (LCV2; a gift from Feng Zhang; Addgene plasmid #52963).

### Cell culture

All cell lines are listed in the **KEY RESOURCES TABLE**.

HEK293T (human kidney epithelial; RRID:CVCL_0063) and A549 (human lung epithelial adenocarcinoma; RRID:CVCL_0023) cell lines were cultured in Dulbecco’s Modified Eagle’s Medium (DMEM; Life Technologies); the KHYG-1 (human leukemic Natural Killer; RRID:CVCL_2976) cell line was cultured in Roswell Park Memorial Institute medium (RPMI-1640; Gibco). Culture media were supplemented with 10% fetal calf serum (FCS; Sigma), 100 U/mL penicillin and 100 µg/mL streptomycin (GIBCO). HEK293T and A549 cells were cultured at 37°C with 10% CO_2_ in a humidified incubator and KHYG-1 cells were cultured at 37°C with 5% CO_2_ in a humidified incubator. Cells were routinely screened for mycoplasma.

### Transient exogenous expression

HEK293T or A549 cells were transfected with FuGENE6 (Promega) or Lipofectamine 2000 (ThermoFisher), respectively, according to the manufacturer’s instructions. 48 h post-transfection cells were lysed directly or re-plated for further experiments.

### Stable exogenous expression

To generate stable cell lines, HEK293T cells were co-transfected with lentiviral constructs (10 µg) and packaging vectors (2.5 μg of pMDL, 5 µg of RSV-REV and 3 μg of VSV)^64^ using Lipofectamine 2000 (Invitrogen) according to the manufacturer’s instructions. Media was replaced 24 h post-transfection and cells were incubated for a further 48 h. Lentivirus was harvested and passed through a 0.45 µm filter prior to incubation with target cells for 24 h followed by recovery in fresh media supplemented with 2.5 µg/mL puromycin. The pFTRE3G DNA constructs required cells to be treated with doxycycline (1 µg/mL) overnight to induce expression.

### CRISPR/Cas9 genome editing

*ARAP2^-/-^* and *SOCS1^-/-^* A549s were generated based on CRISPR/Cas9 protocol described previously (Baker and Masters, 2018) and as detailed above in “**Stable exogenous expression”**. Gene disruption was confirmed by Next Generation Sequencing (NGS). *ARAP2^-/-^* HEK293Ts were with two guide RNAs (gRNAs) targeting exon 2 were synthesized by Synthego. Synthego gRNAs were combined with recombinant Cas9 protein for nucleofection of HEK293T cells, in accordance with the manufacturer’s instructions. Cells were cloned by serial dilution and gene disruption was confirmed by Sanger Sequencing.

### siRNA depletion

All siRNA sequences are listed in the **KEY RESOURCES TABLE**.

A549 cells were reverse transfected with 100 nM siRNA using Lipofectamine RNAi Max transfection reagent (Invitrogen) according to the manufacturer’s instructions. In all experiments, a single scramble siRNA and three ARAP2-targeting siRNAs (in combination) were used. When multiple siRNAs were combined, each siRNA was used at 25 nM and control non-targeting siRNA was used as required to equalize siRNA amounts. Cells were transfected for 48 h prior to re-plating for indicated experimental conditions.

### Cell stimulation

All stimuli and inhibitors are listed in the **KEY RESOURCES TABLE**.

Where appropriate, cells were treated with the following inhibitors or ligands. MG132 was dissolved in DMSO as a 40 mM stock and used at a final concentration of 10 μM for 4 h. Sodium orthovanadate stock solution was prepared as previously described^65^ and activated to pervanadate with hydrogen peroxide for 10 min on ice prior to use^66^. Cells were treated with 50 μM pervanadate for 30 min. Recombinant cytokines were used at a final concentration of 100 ng/mL for times indicated in the figure legends.

### Immunoblots

All antibodies are listed in the **KEY RESOURCES TABLE** and all lysis buffer recipes are detailed in **Table 2**.

**Table 2.**
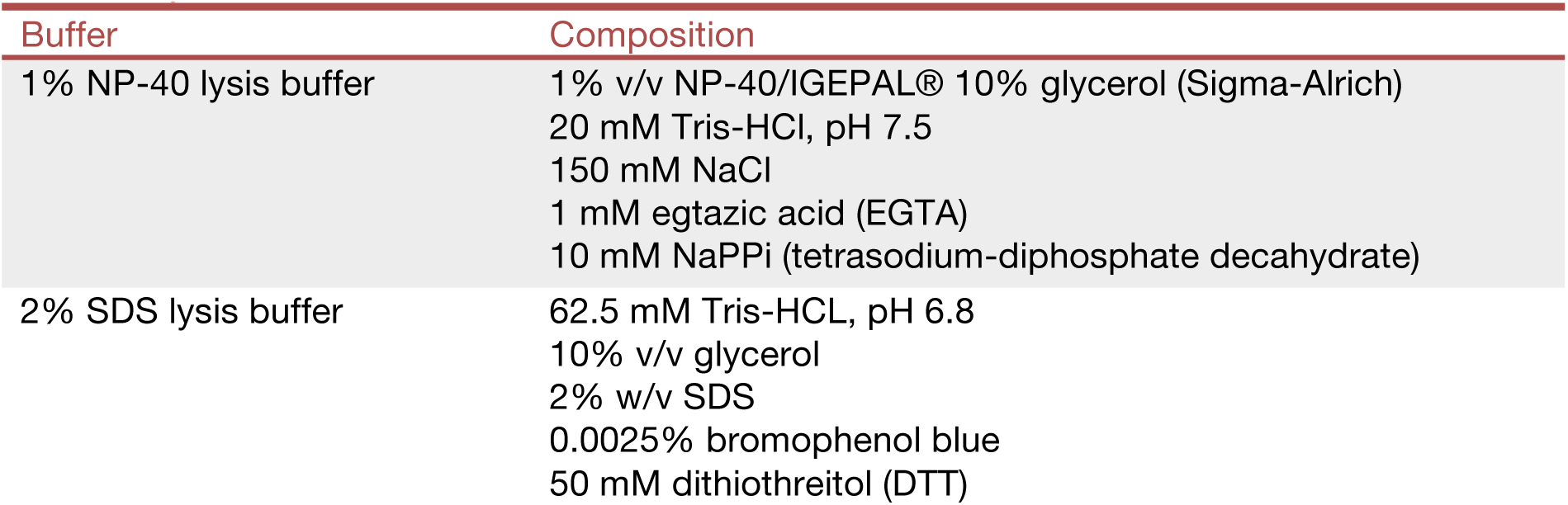
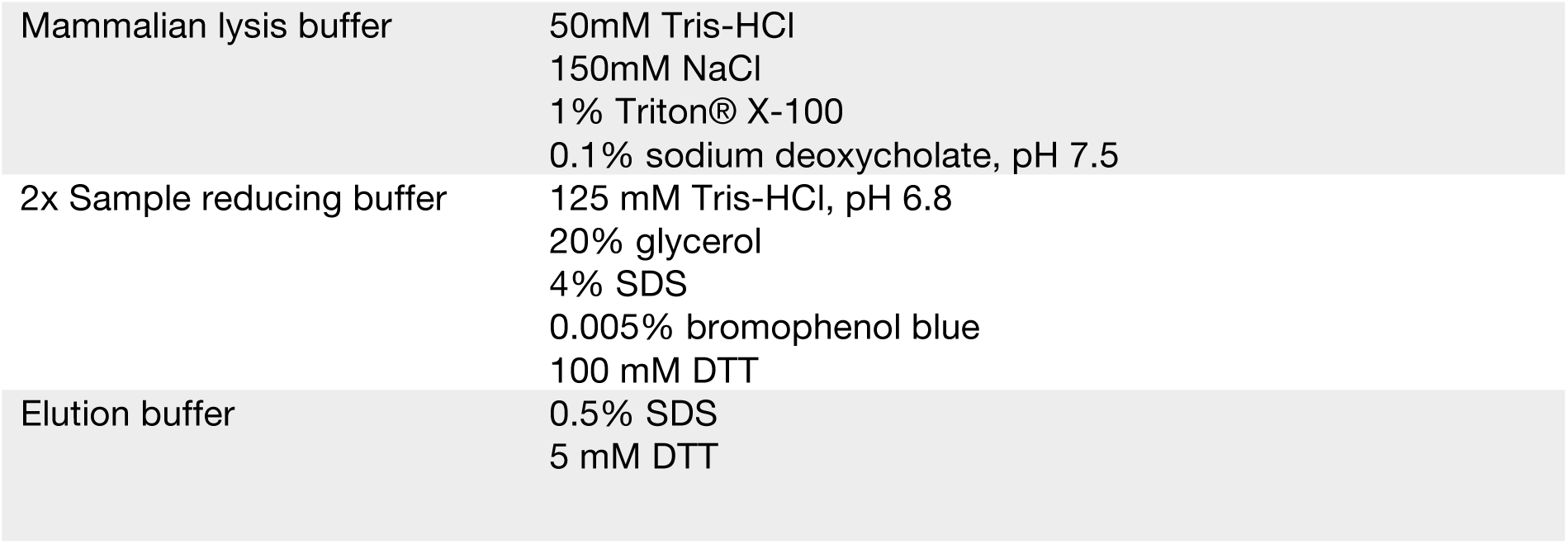
Lysis buffers.

Cells were lysed in 2% sodium dodecyl sulfate (SDS) lysis buffer with β-mercaptoethanol (143 mM). Cell lysates were centrifuged through a 20 µL filter tip to shred DNA. Proteins were separated by sodium dodecyl sulphate-polyacrylamide gel electrophoresis (SDS-PAGE) using NuPAGE® Bis-Tris 4-12 % gels (Invitrogen) in MES buffer [50 mM compound 2-ethanesulfonic acid (MES), pH 7.3] under reducing conditions and electrophoretically transferred to nitrocellulose membranes (Amersham). Ponceau S staining was performed routinely to evaluate protein loading accuracy. Membranes were blocked with 10% skim milk (w/v; Devondale) in Tris-buffered saline (TBS) containing 0.1% Tween 20 (TBS-T) for 20 min. Membranes were probed with primary antibodies overnight at 4°C [diluted in 3% v/v bovine serum albumin (BSA) TBS-T with 0.01% sodium azide at 1:1000]. Secondary antibodies conjugated to horseradish peroxidase (HRP; diluted in 1% w/v BSA TBS-T at 1:10 000) were incubated with membranes for 1 h at room temperature. Membranes were washed with TBS-T between antibody incubations and prior to visualization. Membranes were developed using enhanced chemiluminescence (Millipore, Bio-Rad) using the ChemiDoc Touch Imaging System (Bio-Rad) and Image Lab software.

### Immunoprecipitation and affinity purification assays

All sepharose beads/resin are listed in the **KEY RESOURCES TABLE** and all lysis buffer recipes are detailed in **Table 2**. For immunoprecipitation (IP) experiments, 8 x 10^6^ cells were lysed on ice in 1% NP-40 lysis buffer supplemented with protease inhibitor cocktail (Calbiochem), 1 mM phenylmethylsulfonyl fluoride (PMSF; Sigma-Aldrich), 5 mM sodium fluoride (Sigma-Aldrich) and 1 mM sodium orthovanadate (Sigma-Aldrich). Cell lysates were pre-cleared using 30 µL protein G sepharose (packed bead volume) at 4 °C on a rotating wheel for 1 h. Pre-clearing beads were removed by centrifuging twice (15,000 *g*, 2 min, 4 °C). Anti-FLAG M2-conjugated beads (15 μL packed bead volume; Sigma) were rotated with cleared cell lysates for 3 h at 4°C. Antibody/sepharose complexes were collected by centrifugation (15,000 *g*, 30 sec, 4 °C), washed with 3 x 1 mL ice cold lysis buffer and mixed with equal volume of 2 x sample reducing buffer.

GST affinity purification assays were performed as above with the following modifications. Lysates were pre-cleared with 5 μg GST and 50 μL glutathione sepharose beads (packed bead volume) rotating for 2 h at 4°C. Cleared lysates were rotated with 5 μg GST-tagged CIS (relative to GST%) for 3 h at 4°C. 15 μL glutathione-sepharose beads (packed bead volume) were added for the final hour at 4°C. For washing, lysates and beads were transferred to Pierce spin columns and washed three times in lysis buffer by centrifugation. Enriched complexes were incubated with 120 µL elution buffer at 60°C for 3 min and eluted by microfuge centrifugation.

For HaloTag® affinity purification assays, 8 x 10^6^ cells were lysed on ice in mammalian lysis buffer supplemented with protease inhibitor cocktail (Promega), 1 mM phenylmethylsulfonyl fluoride (PMSF; Sigma-Aldrich), 5 mM sodium fluoride (Sigma-Aldrich) and 1 mM sodium orthovanadate (Sigma-Aldrich). Cell lysates were pre-cleared using Protein G Sepharose at 4 °C on a rotating wheel for 1 h. Pre-clearing beads were removed by centrifuging twice (15,000 *g*, 2 min, 4 °C). HaloLink® resin (25 μL packed bead volume; Promega) were rotated with cleared cell lysates for 4 h at 4°C. Resin complexes were collected by centrifugation (800 *g*, 2 min, 4 °C), washed with 5 x 1 mL ice cold TBS and mixed with 40 µL 2 x sample reducing buffer.

### Mass spectrometry – Trypsin digestion

Eluates from the GST pulldowns (*n* = 3 per group) and lysate input were prepared for mass spectrometry analysis using the FASP protein digestion method as previously described^67^ with the following modifications. Protein material was reduced with Tris-(2-carboxyethyl)phosphine (TCEP; 10 mM). Eluates were digested with sequence-grade modified Trypsin Gold (Promega; 1 μg) in 50 mM ammonium bicarbonate (NH_4_HCO_3_) and incubated overnight at 37 °C. Peptides were then eluted with 50 mM NH_4_HCO_3_ in two 40 μL sequential washes and acidified with 1% formic acid (final concentration). The peptides were then lyophilised to dryness using a CentriVap (Labconco) prior to reconstituting in Buffer A (0.1% FA/2% ACN) ready for mass spectrometry analysis.

### Mass Spectrometry - Data analysis

Peptides were separated by reverse-phase chromatography on a 1.9 μm C18 fused silica column (I.D. 75 μm, O.D. 360 μm x 25 cm length) packed into an emitter tip (Ion Opticks, Australia), using a nano-flow HPLC (M-class, Waters). The HPLC was coupled to an Impact II UHR-QqTOF mass spectrometer (Bruker, Bremen, Germany) using a CaptiveSpray source and nanoBooster at 0.20 Bar using acetonitrile. Peptides were loaded directly onto the column at a constant flow rate of 400 nL/min with buffer A (99.9% Milli-Q water, 0.1% formic acid) and eluted with a 90 min linear gradient from 2 to 34% buffer B (99.9% acetonitrile, 0.1% formic acid). Mass spectra were acquired in a data-dependent manner including an automatic switch between MS and MS/MS scans using a 1.5 second duty cycle and 4 Hz MS1 spectra rate followed by MS/MS scans at 8-20 Hz dependent on precursor intensity for the remainder of the cycle. MS spectra were acquired between a mass range of 200–2000 m/z. Peptide fragmentation was performed using collision-induced dissociation (CID).

Raw files consisting of high-resolution MS/MS spectra were processed with MaxQuant (version 1.5.8.3) for feature detection and protein identification using the Andromeda search engine^68^. Extracted peak lists were searched against the *Homo sapiens* database (UniProt, October 2016), as well as a separate reverse decoy database to empirically assess the false discovery rate (FDR) using strict trypsin specificity allowing up to 2 missed cleavages. The minimum required peptide length was set to 7 amino acids. In the main search, precursor mass tolerance was 0.006Da and fragment mass tolerance was 40ppm. The search included variable modifications of oxidation (methionine), amino-terminal acetylation, the addition of pyroglutamate (at N-termini of glutamate and glutamine), phosphorylation (serine, threonine, tyrosine) and a fixed modification of carbamidomethyl (cysteine). The “match between runs” option in MaxQuant was used to transfer identifications made between runs within a group of samples on the basis of matching precursors with high mass accuracy^69^. PSM and protein identifications were filtered using a target-decoy approach at an FDR of 1%. Statistically relevant protein expression changes between the GST control and GST-CISwere identified using a custom in-house designed pipeline as previously described^70^ where quantitation was performed at the peptide level. Probability values were corrected for multiple testing using Benjamini–Hochberg method. Cut-off lines with the function y = -log_10_(0.05)+c/(x-x_0_) ^71^ were introduced to identify significantly enriched proteins. c was set to 0.2 while x_0_ was set to 1, representing proteins with a twofold (log2 protein ratios of 1 or more) or fourfold (log2 protein ratio of 2) change in protein expression, respectively.

The mass spectrometry proteomics data have been deposited to the ProteomeXchange Consortium via the PRIDE partner repository^72^ with the dataset identifier PXD060300.

### Immunofluorescence and image analysis

4 x 10^4^ A549 cells were plated onto chambered coverslips (IBIDI) prior to stimulation with IFNγ for 30 min. A549 cells were fixed in Cytofix (BD Biosciences) and blocked and permeabilized in PBS containing 5% (v/v) horse serum and 0.2% (v/v) Triton X-100 (blocking solution) for 1 h at RT. Cells were incubated with anti-STAT1 antibody (1:200 in blocking solution) overnight at RT, washed 3-5x in PBS and stained with an anti-rabbit Ig AlexaFluor647 secondary antibody (1:400 in blocking solution) for 2 h at RT. Cells were washed 3-5x in PBS and stored in FluoroShield DAPI mounting media (Abcam) prior to analysis with a Zeiss LSM 880 confocal microscope. The nucleus of the cell was defined by DAPI-positive pixels and nuclear STAT1 was quantified from at least 50 individual cells per condition using the CellProfiler program.

### Proximity ligation assays

Cells were fixed, permeabilized and stained with primary antibodies as detailed above. Proximity ligations was performed using the Duolink® Proximity Ligation Assay kit (Merck) according to manufacturer’s instructions. Analysis was performed with a Zeiss LSM 880 confocal microscope.

### Competitive surface plasmon resonance (SPR) assays

Competitor phosphopeptides were N-terminally amidated and C-terminally acetylated. Peptides bound to the chip were N-terminally biotinylated and C-terminally acetylated. All peptides are listed in the **KEY RESOURCES TABLE**. All SPR experiments were performed on a Biacore 4000 (GE Healthcare) as previously described^58^. Biotin-phosphopeptides were immobilized to a streptavidin-coated SA Biosensor Chip (GE Healthcare) according to the manufacturer’s instructions. 100 nM of recombinant CIS, SOCS1, SOCS2 or SOCS3 in complex with Elongins BC, was mixed with serially diluted competitor ARAP2 phosphopeptides (10 μM, 3 μM, 1 μM, 0.3 μM, 0.1 μM and 0 μM) and flowed over immobilized phosphopeptides (biotin-ARAP2 pY415 for CIS binding, biotin-JAK3 pY980, 981 for SOCS1 and SOCS2 binding, and biotin-gp130 pY758 for SOCS3 binding). Binding was measured using a multiple-cycle method with 60 sec injection, 240 sec association and 120 sec disassociation at a flow rate of 30 μL/min. Response units were expressed as a percentage of maximal binding in the absence of competitor and plotted against the concentration of competitor peptide. IC_50_ values were determined using a non-linear regression and are derived from three independent experiments.

### Luciferase assay

HEK293T cells were transiently co-transfected in technical triplicate with GAS- or ISRE-Firefly luciferase reporter constructs together with a constitutive Renilla-luciferase construct (Cignal Reporter Assay Kit; QIAGEN) using FuGENE6 as per the manufacturer’s instructions. 24 h post-transfection, cells were treated with IFNγ or IFNα for 16 h and reporter activity measured with the Dual-Luciferase Reporter Assay System (Promega) according to the manufacturer’s instructions. Firefly luciferase activity was normalized to Renilla luciferase activity. Luminescence was captured on a ClarioSTAR plate reader (BMG LabTech).

### Mice

The *Arap2^-/-^* and *Arap2^Y413F^* mice were generated by the MAGEC laboratory (WEHI) on a C57BL/6J background. 20 ng/μL of Cas9 mRNA, 10 ng/μL of sgRNA (GTCAGAATACTCCACAGTAG) and 40 ng/μL of oligo donor (ATTTGGATACCCCGCGAGGATAAAAATAACCTTGCACAGGATTCGGCTTCTGAGTCAGAATttagCACcGTgGAGGAATG CTTCCAGAGTTTAAGAAGAAAAAATTCTAAAGCGTCCAAGTCTAGGACTCAAAAAG) were injected into the cytoplasm of fertilized one-cell stage embryos generated from wild type C57BL/6J breeders. 24 h later, two-cell stage embryos were transferred into pseudo-pregnant recipient female mice. Viable offspring were genotyped by next-generation sequencing.

### Immunophenotyping of *Arap2^Y413F^* and *Arap2^-/-^* mice by flow cytometry

All antibodies are listed in the **KEY RESOURCES TABLE** and all buffer recipes are detailed in **Table 3**.

**Table 3.**
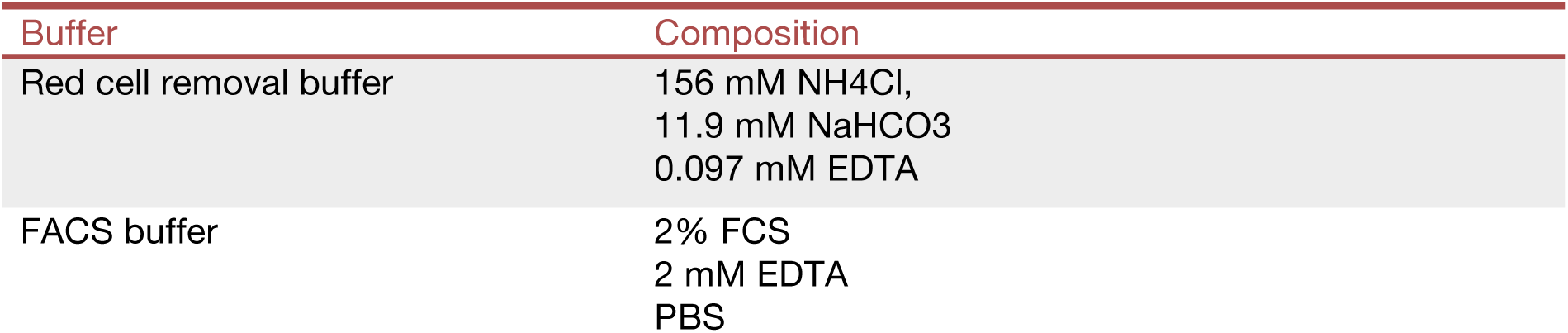
Flow cytometry buffers.

| Buffer | Composition |
| --- | --- |
| Red cell removal buffer | 156 mM NH<sub>4</sub>Cl,<br>11.9 mM NaHCO<sub>3</sub><br>0.097 mM EDTA |
| FACS buffer | 2% FCS<br>2 mM EDTA<br>PBS |

Thymus, spleens and mesenteric lymph nodes were collected from 7-10 week-old female mice (n = 4 per genotype). Single cell suspensions were generated by mashing organs through a 70 μM cell strainer followed by treatment with red cell removal buffer. Cells were FC blocked (FcR Blocking Reagent, mouse, Miltenyi) for 10 min and stained for cell surface markers of interest for 30 min at 4 °C in the dark. Cells were washed twice with FACS buffer, resuspended in FACS buffer with 1 μg/mL propidium iodide (PI) and analyzed using a BD Fortessa 1 flow cytometer. PI exclusion analysis for each sample was performed with 10,000 single cell, non-debris events using a FSC-A versus PI FACS plot. Data analysis was performed using FlowJo software (Version 10), gating on single and viable, CD45^+^CD3^+^ cells.

### Influenza infection

Male mice (6-12 weeks old) were lightly anaesthetized by inhalation of methoxyflurane and infected intranasally (i.n.) with 40 plaque-forming units (PFU) of PR8 (A/Puerto Rico/8/1934) influenza virus. Weights were monitored daily for 10 days. Mice were humanely euthanized if they lost 20% of their initial body weight during the experiment.

### Flow cytometry of influenza-infected lungs

Lungs were collected from male mice (n = 3-4 per genotype). Single cell suspensions were generated by digesting lungs in collagenase IV (Gibco) and DNAse I (ThermoScientific) followed by mashing organs through a 70 μM cell strainer. Where indicated, cells were stained with TMR (20 nM) in RPMI 40 supplemented with 10% heat-inactivated FCS for 2 h at 37°C, prior to treatment with red cell removal buffer. Cells were stained with fixable viability dye for 15 min, FC blocked (FcR Blocking Reagent, mouse, Miltenyi) for 10 min and stained for cell surface markers of interest for 30 min at 4 °C in the dark. Cells were washed twice with FACS buffer and analyzed using an Aurora flow cytometer. Fixable viability dye exclusion analysis for each sample was performed with 10,000 single cell, non-debris events using an FSC-A versus viability FACS plot. Data analysis was performed using FlowJo software (Version 10), gating on single and viable cells. If less than 107 cells were recovered post-processing, mice were removed from data analysis.

### Cytokine analysis and plaque assays of influenza-infected lungs

PR8-infected mice (n=8-10 per genotype) were humanely euthanized two or six days post-infection. Lungs were harvested and homogenized in 1 mL DMEM using the TissueLyser II (Qiagen), centrifuged at *15000 g* for 15 min (4 °C) and the supernatant harvested. Cytokine levels were analyzed in technical duplicates using the BioPlex Pro Assay (BioRad), according to the manufacturer’s instructions. Samples were assayed in technical duplicates. Titers of infectious virus in the tissue homogenates were determined by standard plaque assay on Madin-Darbin Canine Kidney (MDCK) cells, as previously described^73^.

### Bioinformatic analysis of single cell RNA sequencing data

Raw gene counts together with cell barcodes and feature (gene) information were downloaded from Gene Expression Omibus (GEO), accession GSE189407. The count data was produced using the mm10 build of the mouse genome. Additional gene annotation was added from the Ensembl database, version 102. The R packages scater, scran, and scuttle (versions 1.32.1, 1.32.0, and 1.14.0 respectively) were utilized for data analysis^74^. It should be noted that the biological conditions in the data - PR8 and Mock lung infections; appear to have been produced separately. For this analysis the data is processed together.

Cells were first filtered using scuttle’s addPerCellQCMetrics and perCellQCFilters functions. A total of 740 cells were filtered of the initial 4,736. Cell composition was then normalized by first clustering together similar cells using scran’s quickCluster function. Scaling factors were calculated for each cluster that were subsequently de-convolved to the cell level for cell specific normalization using scuttle’s computePooledFactors function.

Following filtering and normalization, the top 10% of highly variable genes were identified using scran’s modelGeneVar function. These genes were then used to perform a principal component analysis (PCA) using scran’s fixedPCA function. The top 50 principal components were retained. A tSNE plot was then generated using scater’s runTSNE function with perplexity set to 20, utilizing the identified highly variable genes and retained principal components. For both the PCA and tSNE calculations, R random seed was set to 100. Cell typing was then performed using the SingleR package version 2.6.0^75^, identifying 20 unique cell types. The Tabula Muris Senis lung samples from the 3m mouse age group were used as a cell reference^76^. These data were downloaded from cellxgene. Finally, data integration is performed to bring together cell types across the biological conditions using the fastMNN method available through the batchelor package version 1.20.0^77^.

All tSNE visualizations were produced using scater’s plotReducedDim function and the ggplot2 software package version 3.5.1.

## QUANTIFICATION AND STATISTICAL ANALYSIS

Each data point from immunofluorescence and luciferase quantification represents a technical replicate, multiple cells from the same cell line treated with the same reagents. Results from technical triplicates are presented as mean and standard deviation (S.D.). The statistical significance of comparisons between two groups was determined using an unpaired *t*-test. Each data point from graphs of mouse experiments represents an independent biological replicate (i.e. different mice). *In vivo* data are pooled from at least two cohorts and are displayed as the mean and standard error of the mean (S.E.M.). Statistical comparisons of different genotypic groups receiving the same treatment were performed using a two-way ANOVA, with a Bonferroni (two genotypes) post hoc correction for multiple comparisons. Paired *t*-tests and two-way ANOVA analyses assumed Gaussian distribution and equal standard deviations between experimental and control groups. All graphical data were prepared and analyzed in GraphPad Prism (Version 9). In all analyses, significance was defined at 0.05, with p > 0.05 (n.s.), p < 0.05 (*), p < 0.01 (**), p < 0.001 (***), p < 0.0001 (****).

## SUPPLEMENTAL INFORMATION

Figures S1–S6

Table S1 List of proteins enriched by GST-CIS/BC in affinity purification mass spectrometry experiment, related to Figure 1

---

**Supplementary Figure 1.**
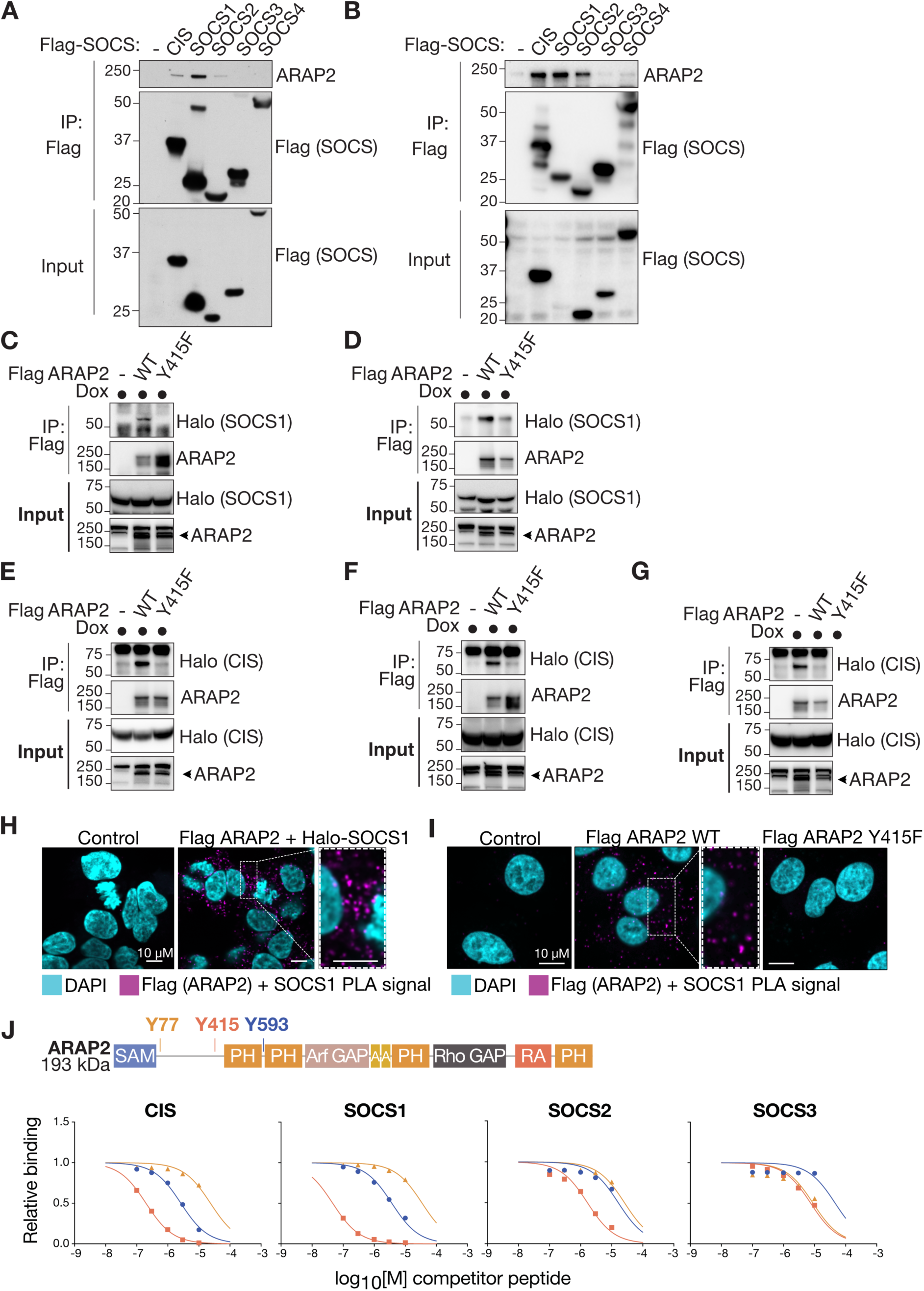
A-B: HEK293T cells were transfected with a panel of Flag-tagged SOCS constructs. Flag-SOCS proteins were immunoprecipitated and enrichment of endogenous ARAP2 analyzed by SDS-PAGE and immunoblot. **C-D:** HEK293T cells stably expressing a doxycycline-inducible HaloTag-SOCS1 construct were transiently transfected with empty pcDNA3.1 vector, Flag-ARAP2 (WT) or Flag-ARAP2 Y415F. Cells were treated with doxycycline overnight to induce SOCS1 expression. Flag-ARAP2 was enriched using anti-Flag antibodies and eluates analyzed by SDS-PAGE and immunoblot. **E-G:** Experiment was performed as in **(C-D)** however, using Halo-CIS-expressing HEK293T cells. **H-I:** Proximity ligation assay (PLA). **H:** HEK293T cells were transiently transfected with Flag ARAP2 and HaloTag-SOCS1. Cells were fixed and stained with anti-SOCS1 antibody and anti-Flag antibody, and DAPI dye (cyan) for confocal analysis using the Zeiss LSM 880 microscope. **I:** A549 cells stably expressing a doxycycline-inducible 3x Flag ARAP2 WT and Y415F construct were treated with doxycycline overnight to induce ARAP2 and stimulated with IFNγ (100 ng/ml) for 6 h to induce endogenous SOCS1 expression. Cells were fixed and stained with anti-SOCS1 antibody and anti-Flag antibody, and DAPI dye (cyan) for confocal analysis using the Zeiss LSM 880 microscope. **J:** ARAP2 domain architecture, highlighting pTyr motifs used in surface plasmon resonance (SPR) assays. Example SPR curves corresponding to data in Table 1 and showing SOCS binding to ARAP2-derived phosphopeptides. Associated with Table 1. Associated with Figure 1.

**Supplementary Figure 2.**
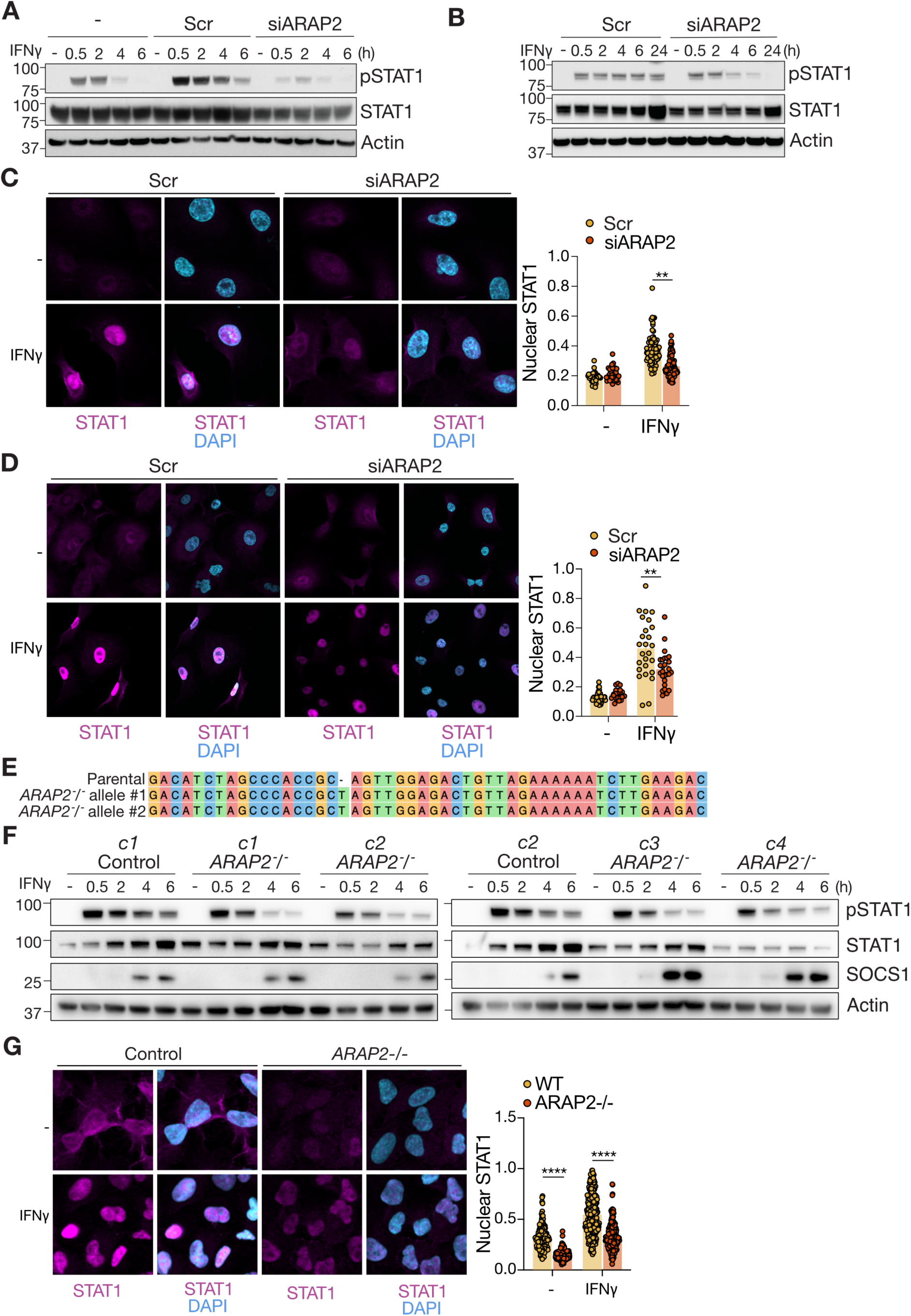
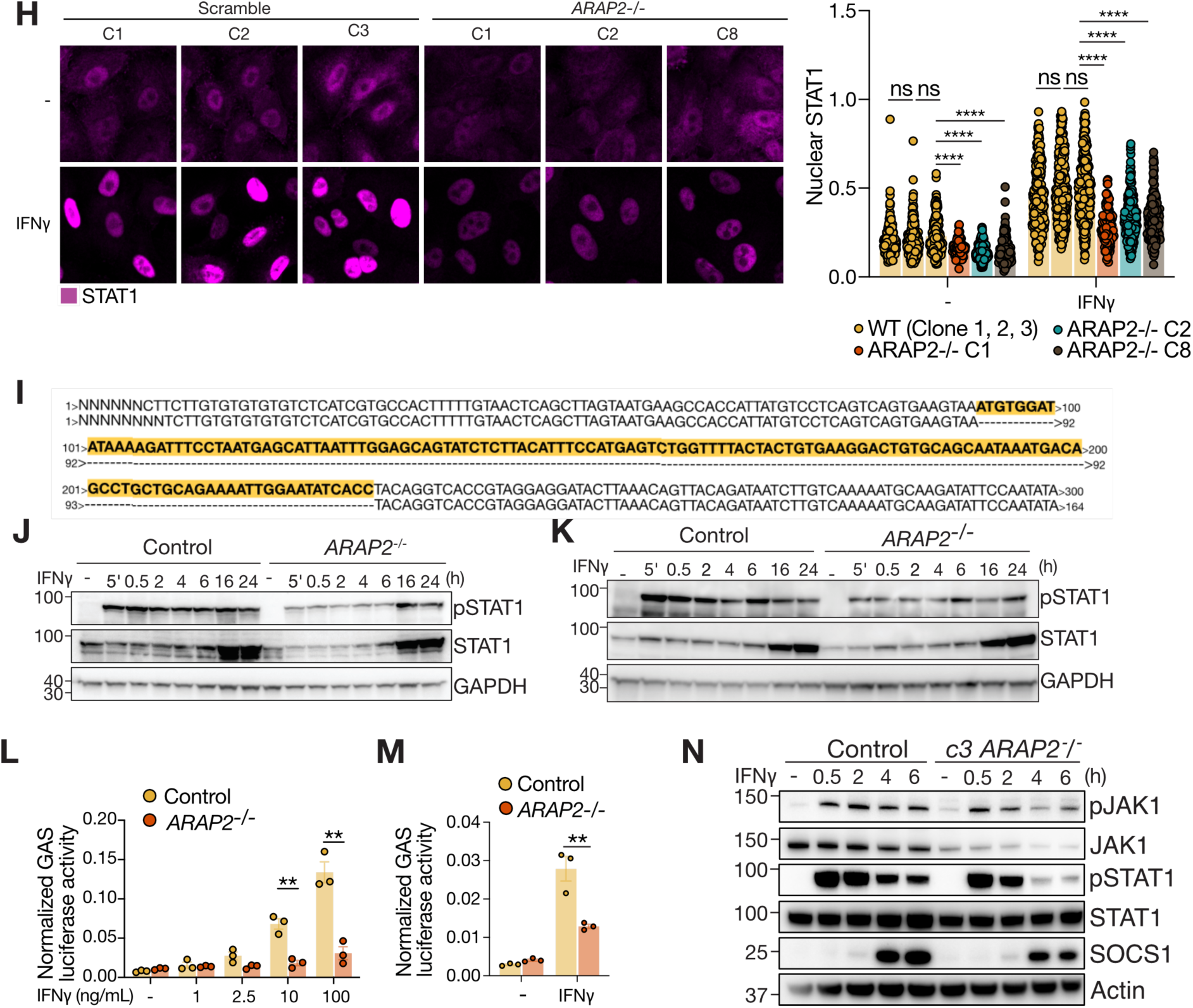
A-B: A549 were transiently transfected with scramble or ARAP2-targeting siRNAs and treated with IFNγ (50 ng/mL) for times indicated. Lysates were analyzed by SDS-PAGE and immunoblot. **C:** Analysis of Next Generation Sequencing. Nucleotide alignment of gDNA generated from parental and *ARAP2^-/-^* A549 cells. The reference parental A549 cells were a pool; the *ARAP2^-/-^*A549 cell line is clonal. **D-E:** A549 were transiently transfected with scramble or ARAP2-targeting siRNAs and treated with 50 ng/mL IFNγ for 30 min prior to fixation. Fixed cells were stained with anti-STAT1 antibody (magenta) and DAPI (cyan) and analyzed using a Zeiss LSM 880 microscope. Quantification of nuclear STAT1. n=50-70 cells. Mean -/+ S.D. Unpaired t-test. **F-G:** WT and *ARAP2^-/-^* A549 cells were treated with IFNγ (100 ng/mL) for the times indicated. Lysates were analyzed by SDS-PAGE and immunoblot. **H-I:** Control and *ARAP2^-/-^* A549 clones were treated with 100 ng/mL IFNγ for 30 min prior to fixation. Fixed cells were stained with anti-STAT1 antibody (magenta) and DAPI (cyan) and analyzed using a Zeiss LSM 880 microscope. Quantification of nuclear STAT1. n=50-70 cells. Mean -/+ S.D. Unpaired t-test. **J-K:** WT and *ARAP2^-/-^* HEK293T cells were transfected with GAS firefly and renilla luciferase constructs in triplicate. IFNγ (100 ng/mL) was added 24 h post-transfection for 16 h before luminescence analysis. Mean -/+ S.D. Unpaired t-test. **L-M:** WT and *ARAP2^-/-^* HEK293T cells were treated with IFNγ (100 ng/mL) for times indicated and lysates were analyzed by SDS-PAGE and immunoblot. **N:** Sanger sequencing from *ARAP2^-/-^* HEK293T clone. The deleted segment is highlighted in yellow. Associated with Figure 2.

**Supplementary Figure 3.**
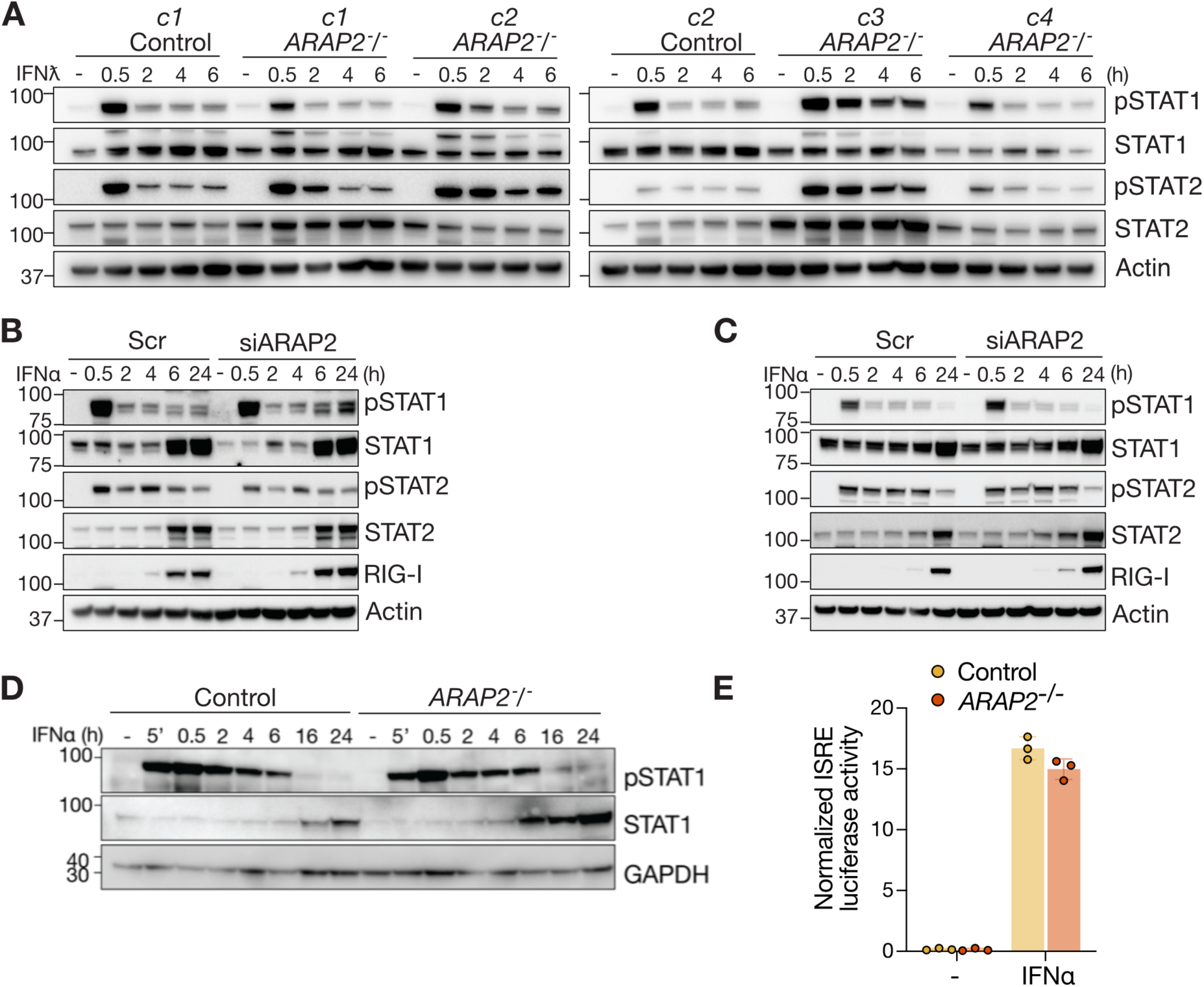
**A:** Clonal scramble sgRNA and *ARAP2^-/-^* A549 cells were stimulated with IFNƛ (100 ng/ml) for times indicated and lysates analyzed by SDS-page and immunoblot. **B-C:** A549 were transiently transfected with scramble or ARAP2-targeting siRNAs and treated with IFNα (50 ng/mL) for times indicated. Lysates were analyzed by SDS-PAGE and immunoblot. **D:** WT and *ARAP2^-/-^* HEK293T cells were treated with IFNα (100 ng/mL) for times indicated and lysates were analyzed by SDS-PAGE and immunoblot. **E:** WT and *ARAP2^-/-^* HEK293T cells were transfected with ISRE firefly reporter and renilla luciferase constructs in triplicate. IFNα (100 ng/mL) was added 24 h post-transfection for 16 h before luminescence analysis. Mean -/+ S.D. Unpaired t-test.

**Supplementary Figure 4.**
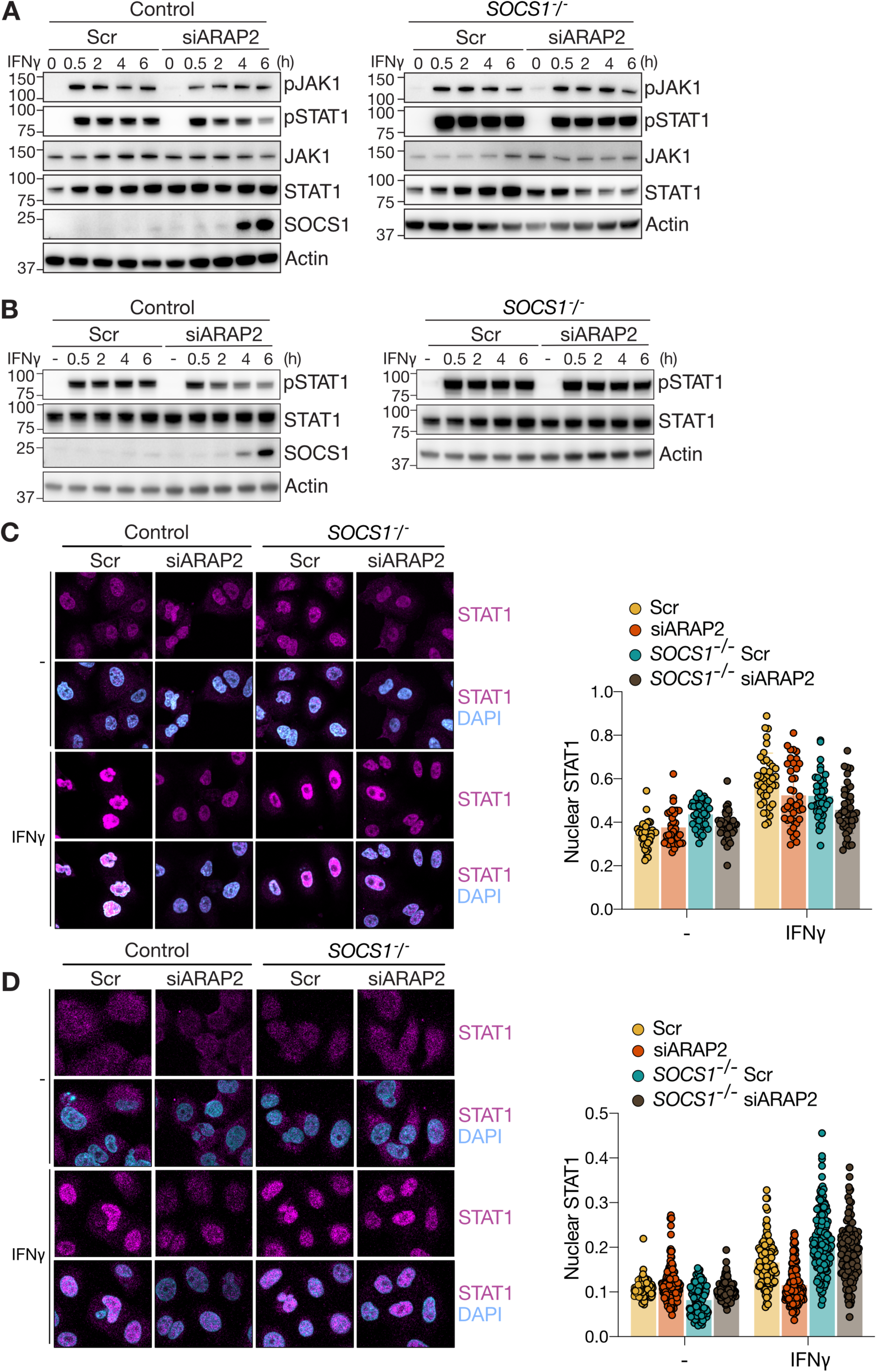
A-B: WT and *SOCS1^-/-^* A549 cells were transfected with scramble or ARAP2-targeting siRNAs and treated with IFNγ (100 ng/mL) for the times indicated. Lysates were analyzed by SDS-PAGE and immunoblot. **C-D:** WT and *SOCS1^-/-^* A549 cells were transfected with scramble or ARAP2-targeting siRNAs and treated with IFNγ (100 ng/mL) for 30 min prior to fixation. Fixed cells were stained with anti-STAT1 antibody (magenta) and DAPI (cyan) and analyzed using a Zeiss LSM 880 microscope. Quantification of nuclear STAT1. n=50-70 cells. Mean -/+ S.D. Unpaired t-test. Associated with Figure 3.

**Supplementary Figure 5.**
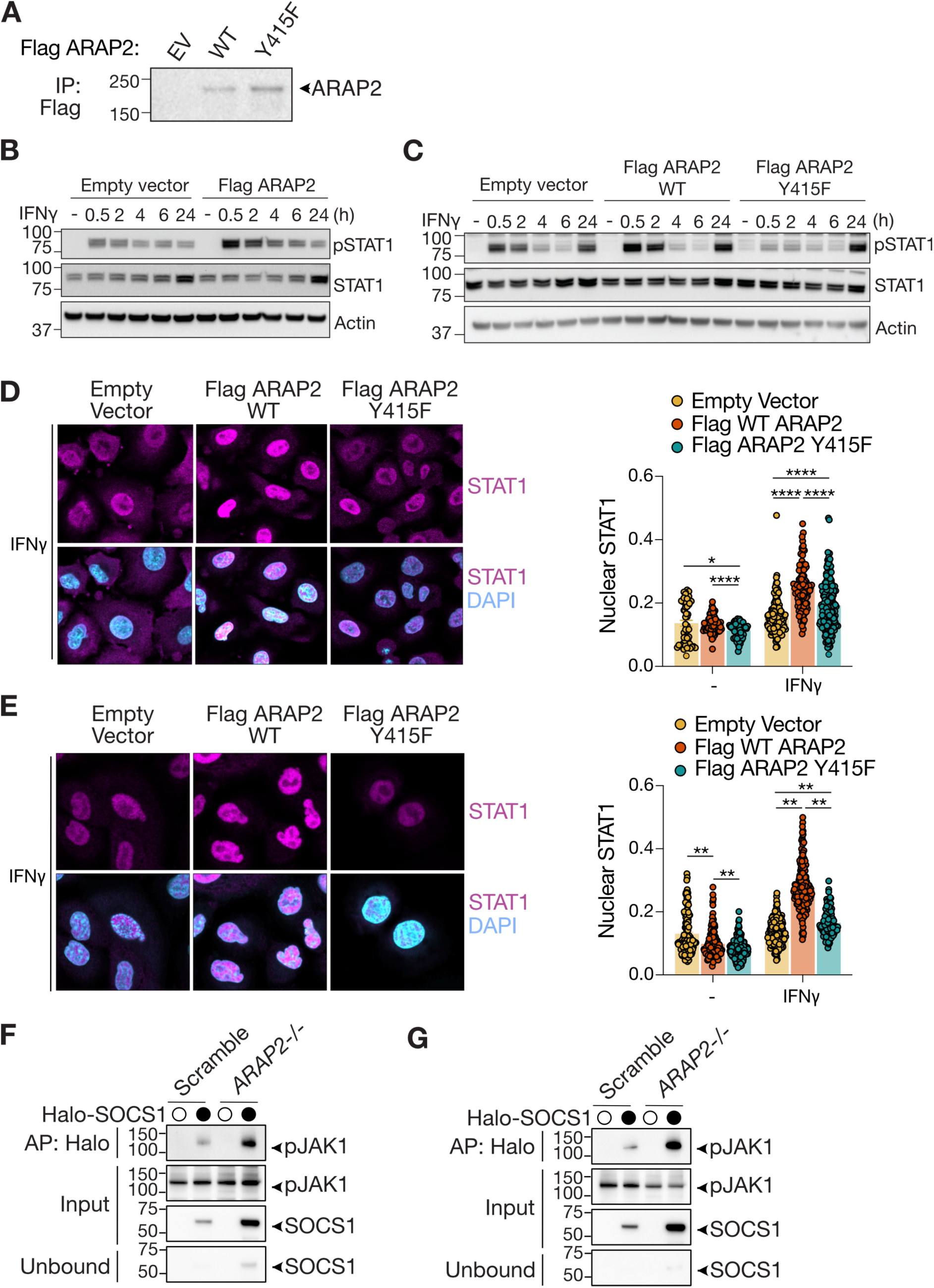
**A:** A549 cells were transiently transfected with empty vector (EV), Flag-ARAP2 WT or Flag-ARAP2-Y415F constructs. 48 h post-transfection, a Flag immunoprecipitation was performed and ARAP2 levels were analyzed by SDS-PAGE and immunoblot. **B:** A549 cells were transiently transfected with pcDNA3.1 empty vector (EV) or Flag-ARAP2 WT. 48 h post-transfection, cells were divided for an IFNγ (100 ng/mL) time course and analyzed by SDS-PAGE and immunoblot. **C:** A549 cells were transiently transfected with pcDNA3.1 empty vector (EV), Flag-ARAP2 WT or Flag-ARAP2 Y415F. 48 h post-transfection, cells were divided for an IFNγ (100 ng/mL) time course and analyzed by SDS-PAGE and immunoblot. **C-D:** Empty vector (EV), Flag-ARAP2 WT and Flag-ARAP2-Y415F transfected A549 cells were stimulated with IFNγ (100 ng/mL) for 30 min prior to fixation and incubation with anti-STAT1 antibody (magenta) and DAPI (cyan) for confocal analysis using the Zeiss LSM 880 microscope. Quantification of nuclear STAT1. n=50-70 cells. Mean -/+ S.D. Unpaired t-test. **E-F:** Control sgRNA and *ARAP2^-/-^* A549s stably expressing a doxycycline inducible construct of Halo-SOCS1 were treated with doxycycline overnight and pervanadate (100 µM) for 30 min prior to lysis. Halo-Flag-SOCS1 was enriched with HaloLink resin and the presence of endogenous pJAK1 were analyzed by SDS-immunoblot. The input and unbound fractions confirm enrichment of SOCS1 during the affinity purification (AP). Associated with Figure 3.

**Supplementary Figure 6.**
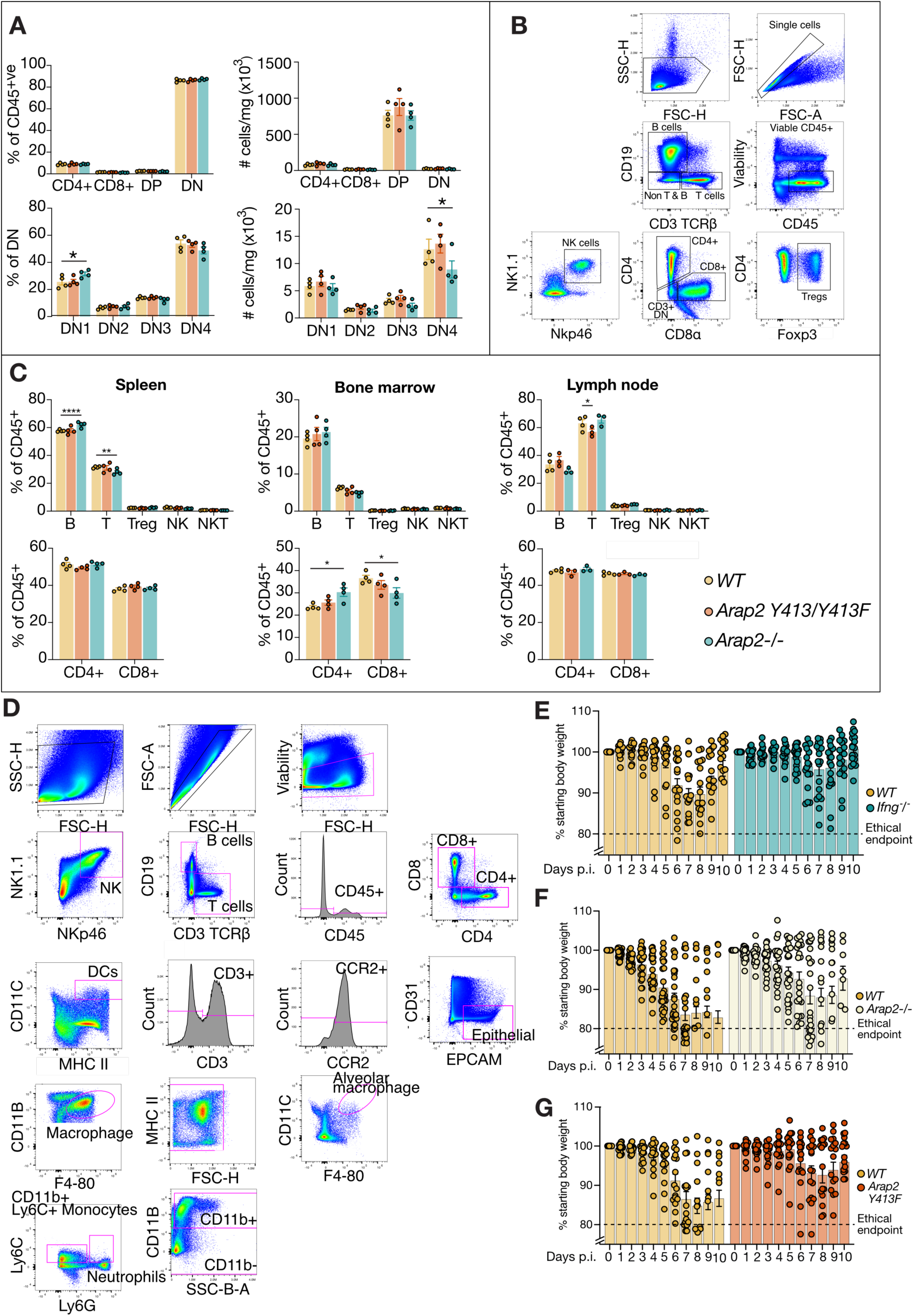
A-B: Immunophenotyping. Thymus, spleen, bone marrow and lymph nodes from WT, *Arap2^Y413F/Y413F^* and *Arap2^-/-^* mice were analyzed by flow cytometry. **A:** Thymus. Cell numbers are per mg thymus. **B:** Example gating strategy. **C:** Spleen, bone marrow and lymph node. **D:** Example gating strategy for flow cytometric analysis of single cell, viable, immune cell populations in lungs from WT, *Halo-Socs1^KI/KI^* and *Arap2^Y413F/Y413F^* mice at day 7 post-infection with PR8 influenza virus. **E-G:** Individual points graphed from Figure 4. Male WT and **(E)** *Ifng^-/-^*, **(F)** *Arap2^-/-^* and **(G)** *Arap2^Y413F/Y413F^* mice were infected intranasally with 40 PFU influenza virus H1N1 PR8 and weight loss monitored for 10 days. Mice that lost more than 20% of their initial body weight were euthanized. Mean ± S.E.M.; n = 18-20. P.I.= post-infection. Associated with Figure 4.

**Supplementary Table 1.**
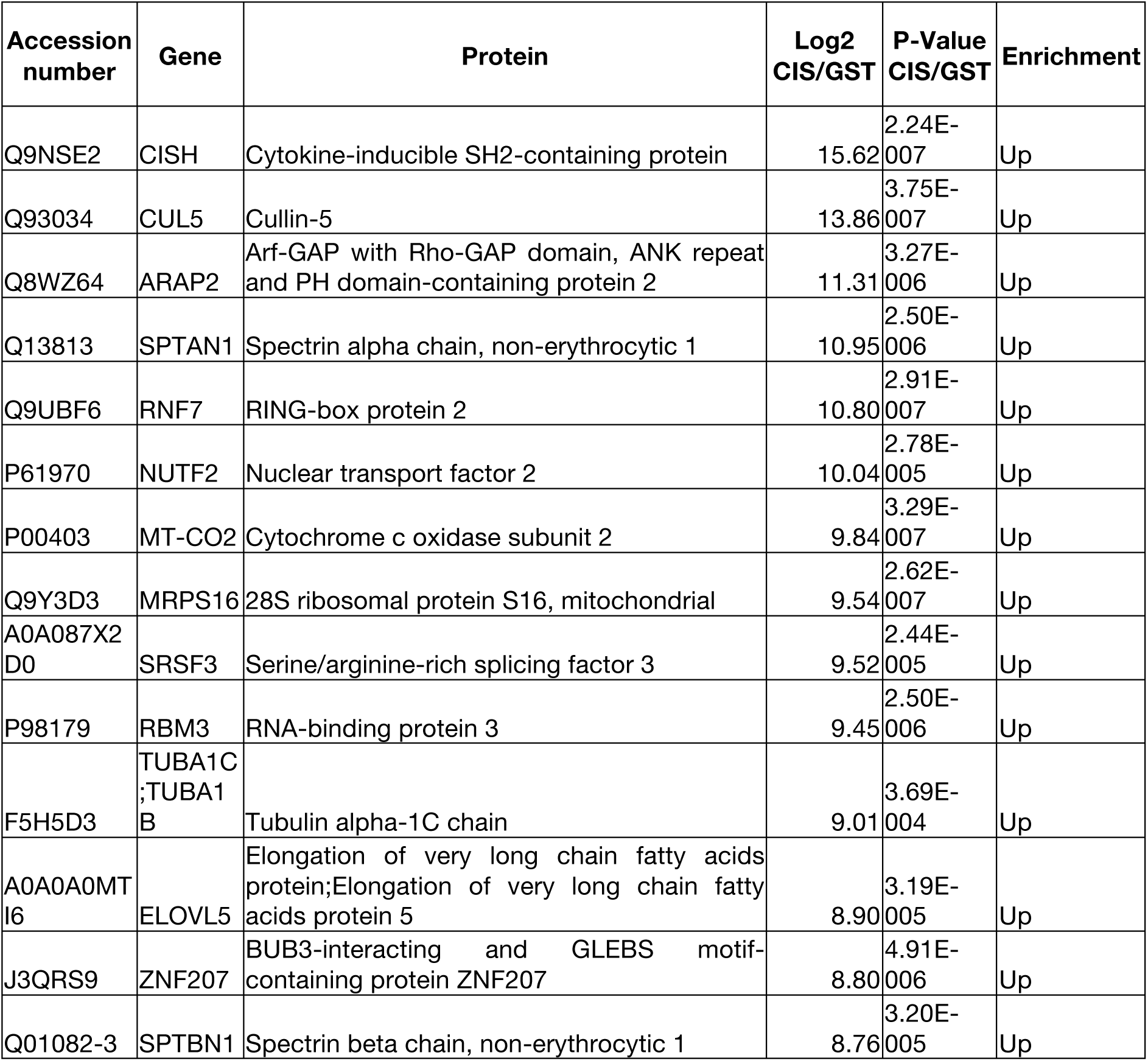
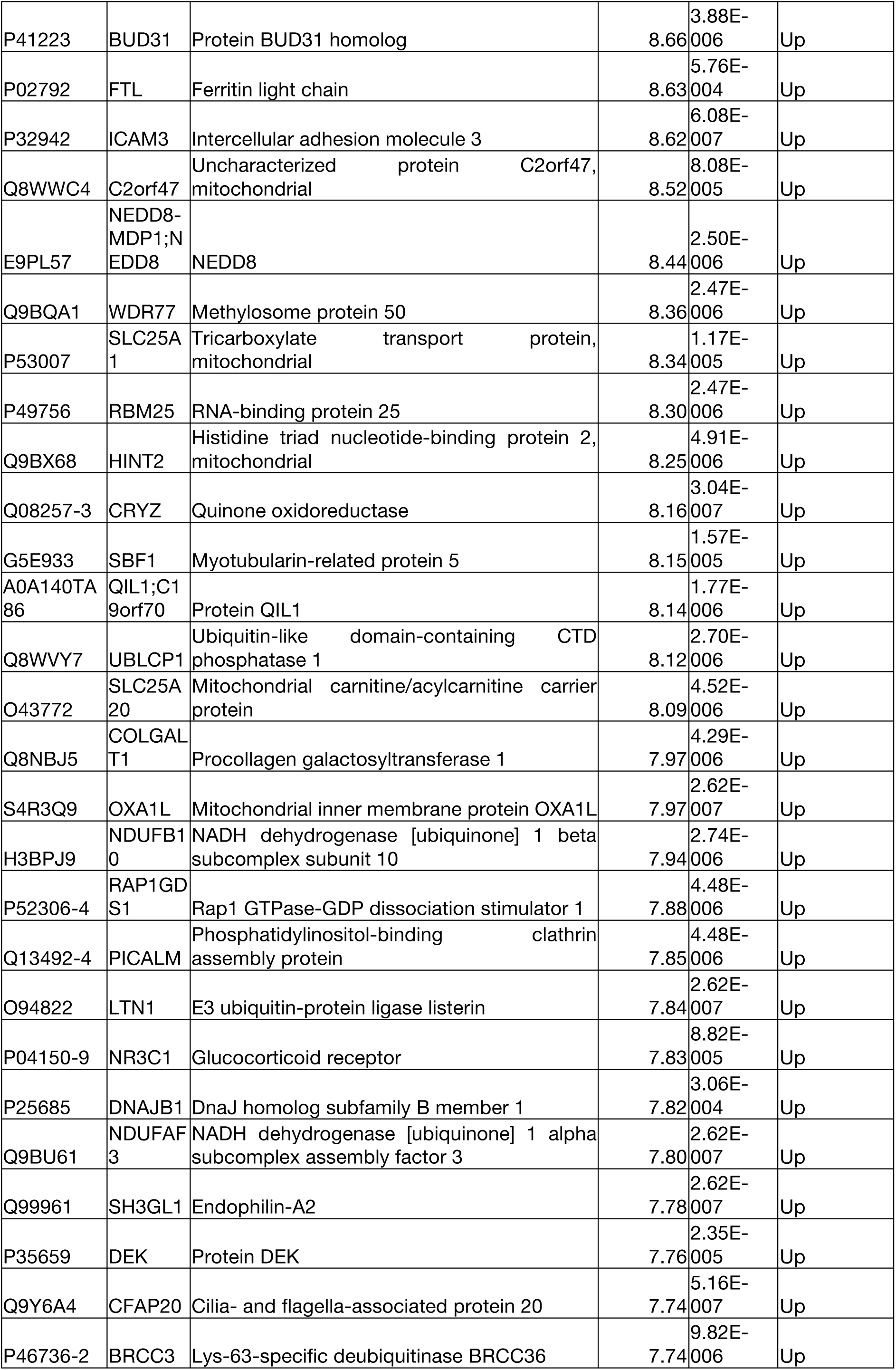
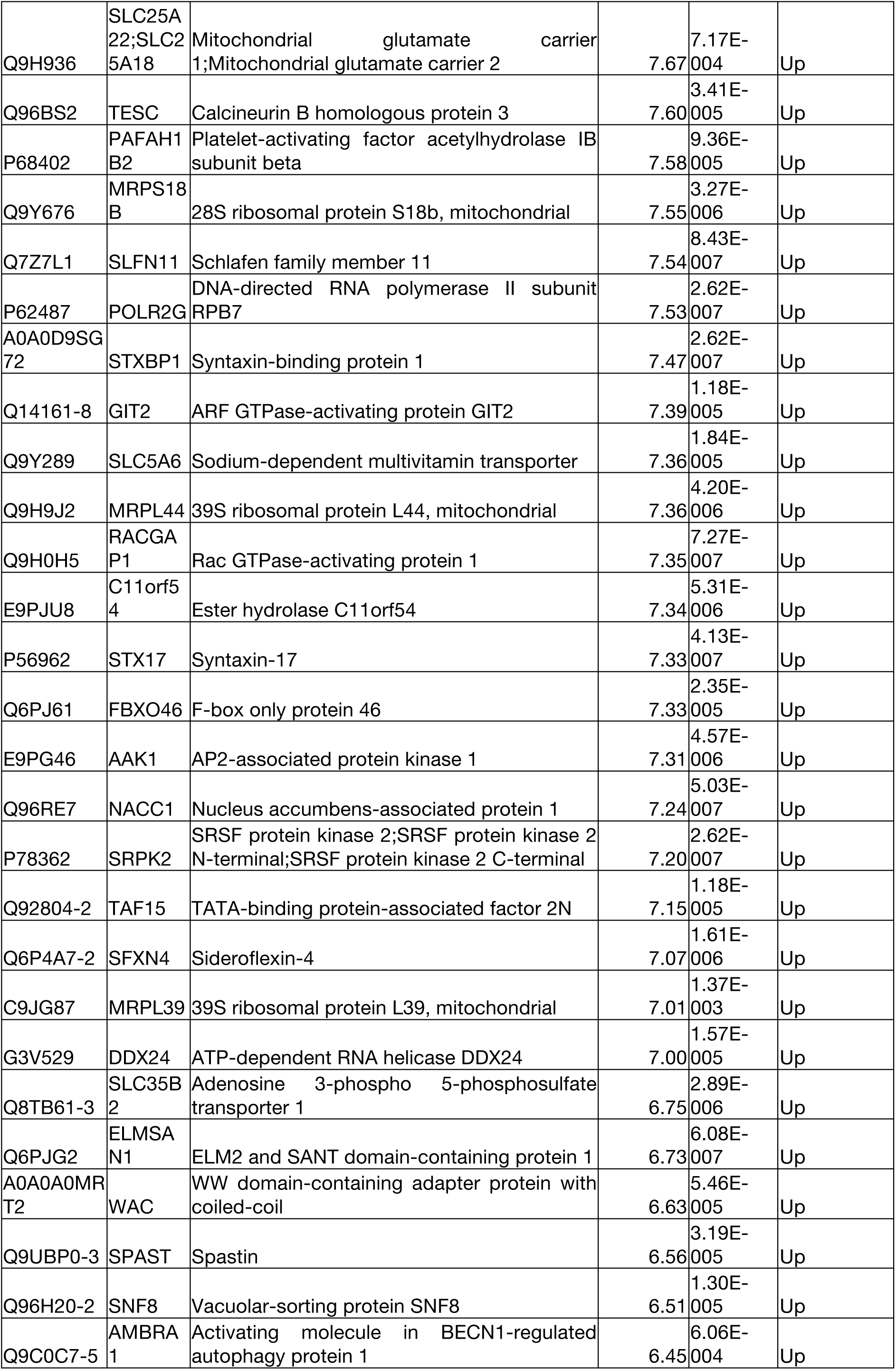
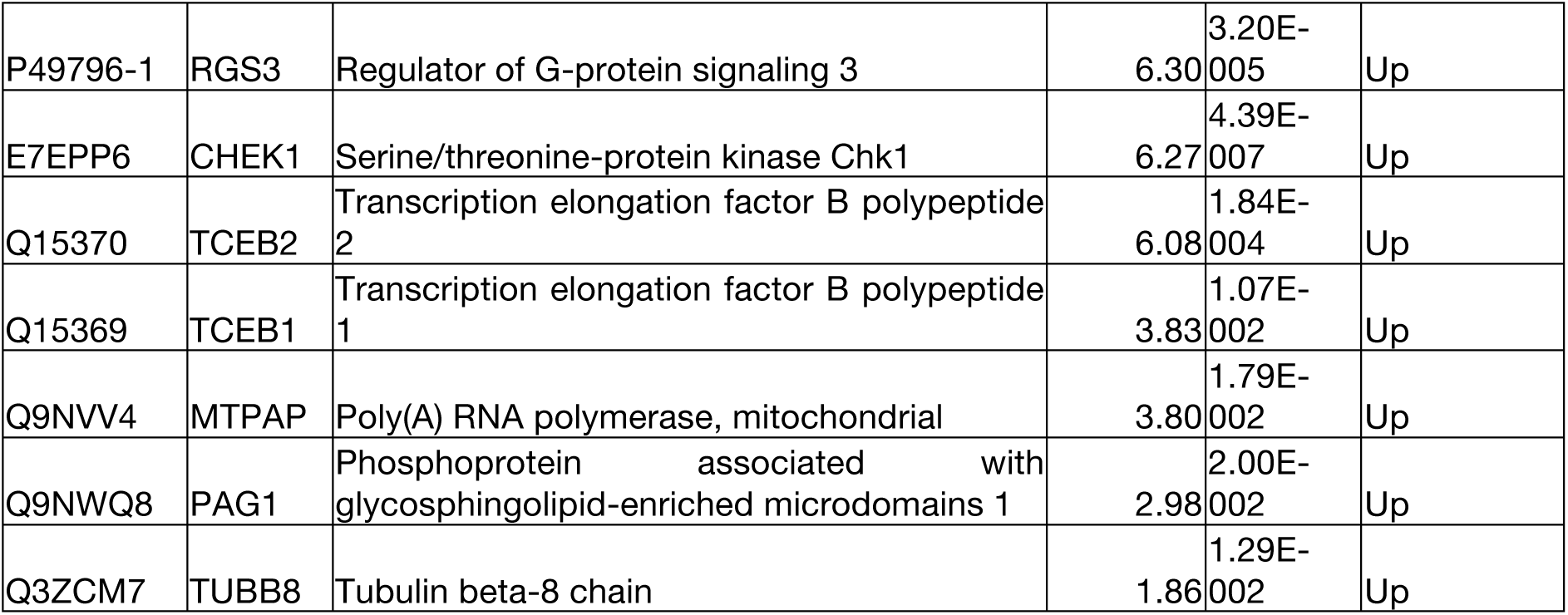
Mass spectrometry data showing proteins differentially enriched with GST-CIS/BC, related to Figure 1A.

